# Evolutionary dynamics of culturally transmitted, fertility-reducing traits

**DOI:** 10.1101/619882

**Authors:** Dominik Wodarz, Shaun Stipp, David Hirshleifer, Natalia L. Komarova

## Abstract

Human populations in many countries have undergone a phase of demographic transition, characterized by a major reduction in fertility at a time of increased resource availability. A key stylized fact is that the reduction in fertility is preceded by a reduction in mortality and a consequent increase in population density. Various theories have been proposed to account for the demographic transition process, including maladaptation, increased parental investment in fewer offspring, and cultural evolution. None of these approaches, including formal cultural evolutionary models of the demographic transitions, have addressed a possible direct causal relationship between a reduction in mortality and the subsequent decline in fertility. We provide mathematical models in which *low mortality* favors the cultural selection of low fertility traits. This occurs because reduced mortality slows turnover in the model, which allows the cultural transmission advantage of low fertility traits to out-race their reproductive disadvantage. For mortality to be a crucial determinant of outcome, a cultural transmission bias is required where slow reproducers exert higher social influence. Computer simulations of our models that allow for exogenous variation in the death rate can reproduce the central features of the demographic transition process, including substantial reductions in fertility within only 1-3 generations. A model assuming continuous evolution of reproduction rates through imitation errors predicts fertility to fall below replacement levels, if death rates are sufficiently low. This can potentially explain the very low preferred family sizes in Western Europe.

## 1. Introduction

In the 19^th^ century, some human populations displayed a demographic transition from relatively high fertility and high mortality towards a greatly reduced fertility and lower mortality [1–4]. This first occurred in more developed parts of the world, such as Europe, the United States, Japan, Australia, and New Zealand, and coincided with an overall increase in resource availability (judged by economic considerations). In Western European countries, fertility declined below replacement levels since the 1970s and 1980s [5,6], and this also applies to preferred family sizes. In German speaking countries the average reported ideal family size has fallen below replacement levels— about 1.7 children [6]. Furthermore, fertility reduction tends to be more pronounced in population segments that are economically advantaged than in poorer segments [1]. This is in contrast to trends observed before these demographic transitions, when wealth was associated with higher fertility [1,7].

A number of theories have been put forward to account for demographic transitions towards reduced fertility [1,8]. According to one line of argument, the transition to reduced fertility may be because of a mismatch between the modern environment and the ancestral one in which humans evolved. Behaviors that were advantageous in the ancestral environment could have become dysfunctional under modern socio-economic conditions, leading to a reduced reproductive output [1]. A second theory holds that the current environment favors the production of few offspring with large parental investment rather than the generation of more offspring with lesser parental investment per child. A third theory is based upon cultural rather than genetic evolution [9,10]. Behavior that leads to reduced fertility in certain influential individuals is copied by others, resulting in a spread of this trait.

A well-developed mathematical theory of the dynamics of cultural transmission [9,11–17] has been applied to the analysis of demographic transitions and the evolution of small family sizes [18–20]. This research has analyzed the spread of cultural traits that affect fertility, survival, or both, and the effects of these traits on the demographic structure of the population. In [19,20], the transition to reduced fertilities has been explained by cultural niche construction. According to this theory, the first trait to spread is one of valuing education, which provides an environment that promotes the spread of a second, fertility-reducing trait. If the trait of valuing education is further associated with reduced mortality of individuals, the model predicts that the decline in fertility is preceded by a reduction in the population death rate, as observed in demographic data. In [18] it was shown that horizontal and oblique transmission can accelerate the spread of the cultural trait, compared to vertical transmission alone. This paper provides a broad analysis and creates a model of cultural transmission of a trait that can affect fertility and/or mortality of individuals. Applications to demographic transitions are described in two contexts: (i) Neolitic demographic transition, where a fertility-increasing trait spreads through the population, is investigated with respect to different transmission modes, and (ii) 19-20 century demographic transition in Europe is modeled by using a trait that simultaneously decreases fertility and increases survival of individuals. A trait is considered where a reduction in fertility is strongly coupled to an increase of survival of individuals.

A key stylized fact about demographic transitions is that the reduction in fertility tends to be preceded by a reduction in the death rate of individuals, and by a consequent temporary population growth phase [4,21,22], presumably a consequence of improved socioeconomic circumstances. This is surprising in the light of evolutionary biology [1,23], because evolution tends to maximize reproductive output, which can generally be increased when resources are more plentiful. Mathematical models of cultural evolutionary processes have so far not directly addressed the reason for the observation that fertility reduction is preceded by mortality reduction. Previously published work linked mortality reduction to other cultural traits, such as education or fertility itself. Here, we add to the existing literature by considering mathematical models of cultural transmission where the population death rate is subject to independent external influences that vary exogenously over time, due to sanitary, medical and technological advances. We investigate how such externally-driven changes in mortality affect the contagion of a fertility-reducing trait. We find that the death rate of individuals is a key parameter for determining whether the cultural spread of a fertility-reducing trait is successful. While the fertility-reducing trait fails to spread at high population death rates, it successfully spreads once the population death rate has fallen below a threshold. For this impact of the population death rate to be observed, the model further requires a cultural transmission bias towards slow reproducers, which can come about by a higher social influence of slowly reproducing individuals. The critical effect of the population death rate on outcome occurs because reduced death rates slow the rate at which early reproducers outrun delayed reproducers in the models, allowing cultural transmission of low fertility traits to outweigh the fitness advantage of fast reproduction. Computer simulations of the demographic transition process show that the empirical stylized characteristics of this process can be captured by our models on realistic time scales. The models further predict that with reduced population death rates, cultural evolutionary processes can result in the eventual decline of fertility below replacement levels. This is relevant for recent trends in Western European and other countries [5,6].

## 2. Concepts and modeling approaches: a roadmap

Cultural transmission dynamics can be complex, and several different mathematical modeling assumptions can be made that can potentially impact results. While simpler models are more tractable analytically, including some more realistic assumptions requires more complicated modeling approaches. Therefore, the paper is structured as follows (Figure 1).

a. We start with the simplest modeling approach that takes into account two distinct populations: fast versus slow reproducers. Moreover, it will be assumed that all individuals mix perfectly with each other, and that logistic growth occurs that is limited by a carrying capacity. This is expressed in terms of ordinary differential equations, and basic insights will be described about the conditions required for slow reproducers to be prevalent.
b. The same kind of dynamics (fast versus slow reproducers) will be re-considered in biologically more complex settings. These include: (i) A spatially explicit model, because the perfect mixing assumption is unrealistic and individuals are more likely to communicate with members of their local community rather than with anyone in the global population. Including spatial restriction has been shown to have significant effects not only in ecological and evolutionary models, but also in models of cultural evolution [24,25]. (ii) An age-structured model where instead of fast and slow reproduction rates, we consider early and late reproducers, because the timing of reproduction can be an important determinant of fertility. (iii) Instead of a fixed carrying capacity, we assume that more room for increased population growth is continuously generated, thus giving rise to an ever-increasing population size, which is more realistic. Using this model, we further show that a demographic transition from higher to lower fertility can occur within realistic time frames. An important conclusion from this section is that central results remain robust irrespective of the modeling approach, thus increasing the confidence in biological / sociological relevance.
c. The longer-term evolution of fertility will be examined. This requires a different approach where the reproduction rate is allowed to continuously evolve, rather than assuming fast versus slow reproducers. The most straightforward way to model this is in terms of an agent-based model, and we will build on the spatial model considered in Bii.

**Figure 1:**
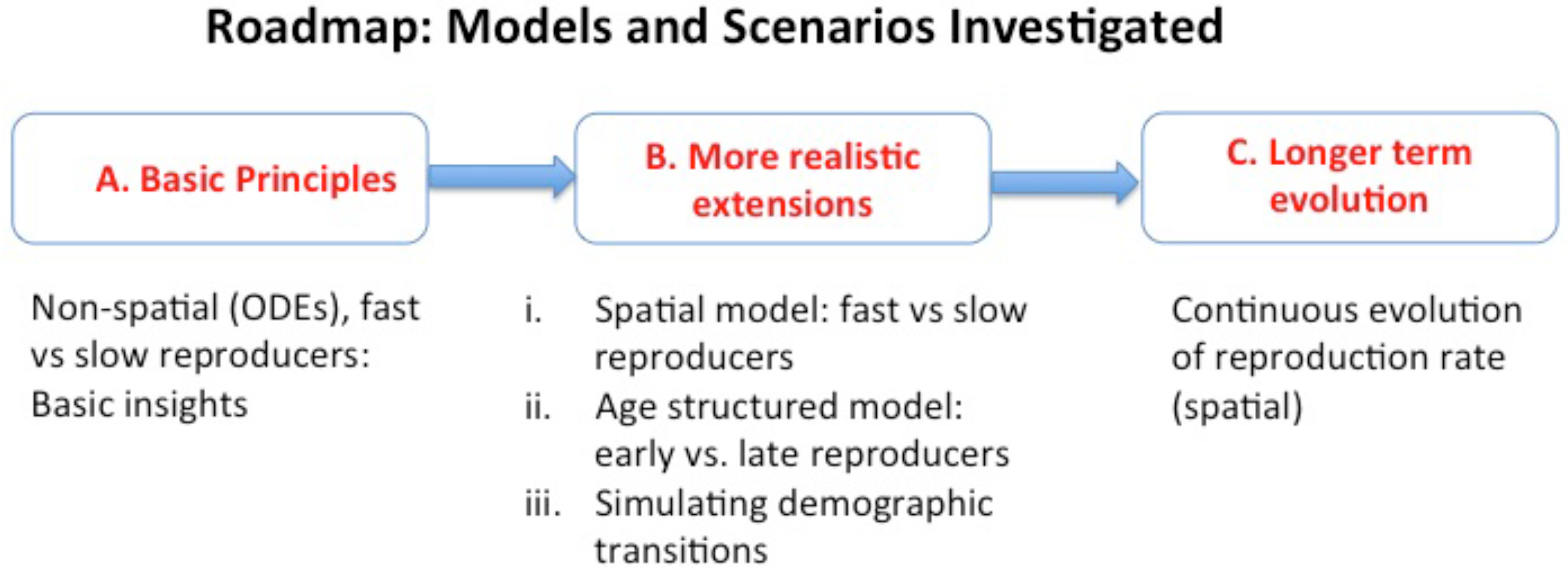
Schematic representation of the structure of the papers and the types of models considered. See text for details.

## 3. Results

### 3A. Fast versus slow reproducers in well-mixed populations

We start the exploration of the evolutionary dynamics of a culturally transmitted, fertility reducing trait by formulating a minimally parameterized model that includes (a) a fertility reducing trait and (b) cultural transmission. We assume that two traits exist in the population. The fast reproduction trait is a default state, and a slow reproductive trait can spread culturally via horizontal or vertical transmission. We will denote the population of the individuals with the fast reproductive trait as x_f_ and the population of the individuals with the slow reproductive trait as xs. The dynamics can be described by a deterministic, non-spatial, asexual model expressed by ODEs:

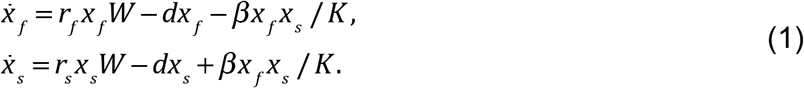

Here, each type reproduces with its own linear reproduction rate, with r_f_>r_s_, and the competition between the two traits is expressed by term W, which for example can take the logistic form,

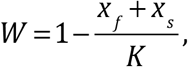

where K denotes the carrying capacity. Both types die with equal rates, d. We assume that there is a probability of switching from one type to the other, which is proportional to the abundance of the individuals of the opposite type. The total rate at which fast reproducing individuals switch to slow reproduction is given by 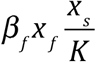, and the total rate at which slow reproducers switch to fast reproduction is given by 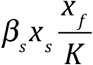. If we assume that β_f_>β_s_, and denote β= β_f_-β_s_, we have the term 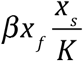 with the negative sign in the equation for x_f_ and the same term with the positive sign in the equation for x_s_. These terms are equivalent in form to infection terms, see equation (1). The main postulates used here are that (a) of the two types of individuals, one grows faster than the other (r_f_>r_s_) and (b) there are more individuals switching from fast reproduction to slow reproduction than the other way around (β>0). The latter modeling choice is motivated by the assumption that slow reproducers tend to channel the resources available to them into accumulation of wealth and/or social status, and thus they may appear as more attractive models for imitation [19].

System (1) has four steady states:

0. The trivial solution, x_f_ = x_s_=0 is unstable as long as r_s_>d and r_f_>d. We will assume that both populations can persist on their own, and the above inequalities hold.
1. Fast reproducers win (that is, the fast reproduction trait spreads through the whole population): *x_f_* = *K*(1 – *d*/*r_f_*), *x_s_* = 0. This solution is stable if

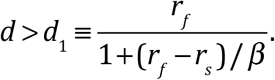
2. Slow reproducers win: *x_f_* = 0, *x_s_* = *K*(1 – *d*/*r_s_*). This solution is stable if 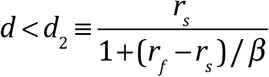. Note that d_2_<d_1_.
3. Coexistence solution, where both traits occur in the population

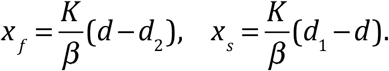 This solution is positive and stable as long as d_2_<d<d_1_.

To summarize these results, we note that the death rate of the individuals, d, controls the outcome of the competition dynamics of the two traits. For high death rates, the fast reproduction trait spreads through the population, and for low death rates, the slow reproduction trait is able to invade and take over. Modifications of the basic model (1) are considered in Section 1.1 of the Supplement, where we study different assumptions on the dynamics of switching type; it is shown that the central results are unchanged. We note that to observe these results in the current setting, the models need to include the assumption of density dependence in the population growth process. They are not observed in models assuming straightforward exponential population growth. Section 3Biii below explores models of unbounded population growth in which the results reported here remain robust.

Before we proceed, it is instructive to interpret the model from the prospective of virus dynamics, by viewing x^(1)^ and x^(2)^ and susceptible and infected individuals respectively. The three nontrivial equilibria are characterized by (1) susceptibles only, (2) infecteds only, and (3) coexistence of both. In order for infection to be able to spread, the basic reproductive ratio, R_0_, has to be larger than 1. In the context of this system, we have 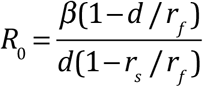.

Decreasing d clearly increases R_0_.

### 3B. Introducing more realism into the model

Because the model explored in the last section contains a number of simplifying assumptions that are known to be inconsistent with reality, it is important to determine whether the results hold robust in more realistic settings. It turns out that central results do remain robust in spatial models, models with age structure, and in models assuming that populations periodically increase their carrying capacity. This is described as follows.

#### (i) Spatial Dynamics

We consider a stochastic agent-based model (ABM) that describes population dynamics on a 2D grid of size *n* × *n*. We will refer to this model as ABM1; compared to the simple ODE model, the present description includes spatial and stochastic effects. As before, the fast reproduction trait is assumed a default state of the agents, and a slow reproductive trait spreads culturally via horizontal or vertical transmission. During each time step (representing a generation), the grid is randomly sampled 2M times, where M is the total number of individuals currently present. When an individual is picked, it attempts to undergo either a birth-death update (including vertical cultural transmission), or a horizontal cultural transmission update. The two types of update are chosen with equal probabilities, such that on average there are M attempts of both types of update during each time step.

If the birth-death update is chosen, the individual can undergo at most one event, as follows. It attempts reproduction with a probability R_f_ or R_s_, depending on whether this is a fast or slow reproducer (here R_f_> R_s_), or dies with a probability D (both populations are assumed to have the same death rate). For a reproductive event, a spot is chosen randomly from the eight nearest neighbors. If that target spot is empty, the offspring is placed there, otherwise, the reproduction event is aborted. We assume that the reproductive strategy of the offspring is the same as that of the parent (that is, the slow reproductive trait is passed on via vertical cultural transmission). These birth-death processes on the grid are characterized by density dependence, and hence the model accounts for competition between slow and fast reproducers. The description above corresponds to infant mortality rising with increased density (crowdedness), because offspring disappear if they do not fall on an empty spot in the grid. Section Biii below explores how such processes can apply to growing human populations.

A cultural update is attempted with probability P_C_, by gathering the information on the reproductive strategy of the individuals’ neighbors, similar to voter models [15,26]. The probability that an agent switches its reproductive strategy is proportional to the weighed fraction of the opposing strategy among neighbors, such that slow reproducers are more influential than fast reproducers. When adding up the number of fast and slow reproducers in the neighborhood, there is a probability Q<1 that a fast reproducer is taken into account, while all slow reproducers are always included, reflecting the preference of switching towards slower reproduction.

When the model is run with only the reproduction and death processes (no non-vertical cultural transmission), then the only outcome is the persistence of the fast-reproducing trait and the competitive exclusion of the slower reproducing one. This is straightforward competition dynamics behavior. If, in contrast, the model is run with only horizontal cultural transmission (no reproduction and death, so that the population is constant), it essentially becomes a voter model, where “slow” and “fast” are different opinions held by individuals in the population. As has been described for such models [15,26], the only eventual outcome is that every individual in the population has the same opinion. Which of the two opinions wins depends on the bias, Q, and on initial frequencies of the opinions in the population.

When we allow for both horizontal transmission and reproduction with vertical transmission, three outcomes are possible (Figure 2): (1) The fast reproduction trait wins and excludes the slow reproduction trait. (2) The slow reproduction trait wins and excludes the fast reproduction trait; and (3) both traits coexist in a long term equilibrium. While this is a true equilibrium in corresponding ODEs (see above), the stochastic nature of the model means that the eventual outcome is always extinction. The coexistence outcome, however, is characterized by a significantly longer time to extinction compared to the exclusion outcomes (compare Figure 2C to 2A & B).

**Figure 2.**
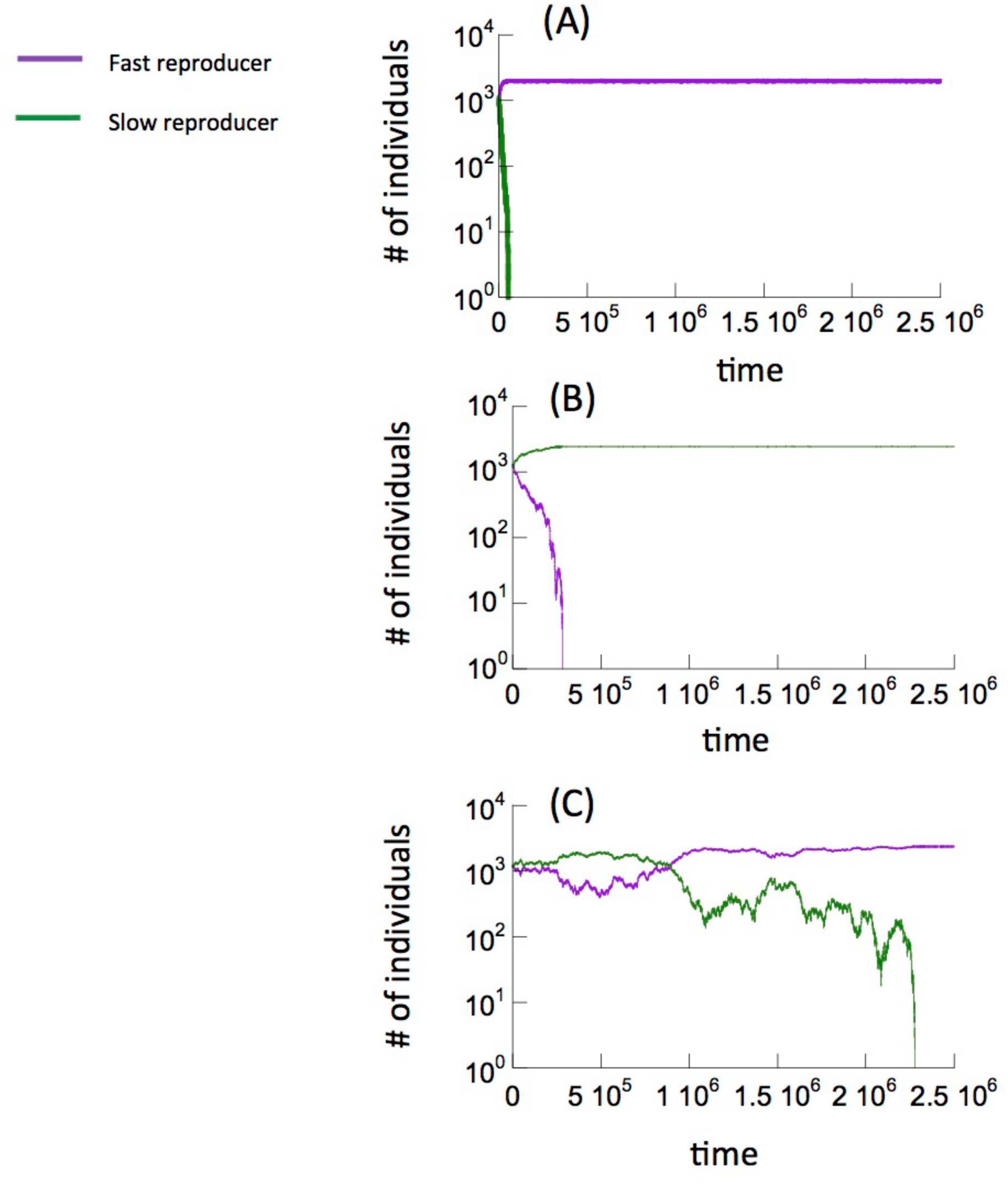
Time series showing the different outcomes according to ABM1. Individual realizations are shown. (A) Higher death rates: the fast-reproducing trait persists and the slow-reproducing trait goes extinct on a short time scale. (B) Lower death rates: the slow-reproducing trait persists and the fast-reproducing trait goes extinct on a short time scale. (C) Intermediate death rates: both fast- and slow-reproducing traits persist for significantly longer time periods. Eventually one trait goes extinct due to the stochastic nature of the simulation. Parameters were chosen as follows. R_f_=0.005; R_s_=0.8R_f_; P_C_=0.0008; Q=0.93. For (A), D=0.001. For (B), D=0.0001. For (C), D=0.00025.

Which outcome is observed depends on the death rate of agents, D, see Figure 3Ai. Each point on this graph depicts the time until one of the traits goes extinct, depending on the death probability, D. The outcomes are color-coded: purple depicts fast reproducers remaining, and green slow reproducers. At higher death rates, the fast reproducers persist and extinction of the slow reproduces occurs at relatively short time scales. At low death rates, the slow reproducers persist and the fast reproducers go extinct on a relatively short time scale. At intermediate death rates, the time to extinction of one of the populations rises sharply, and either population has a chance to go extinct first. This corresponds to the coexistence regime. Therefore, lower death rates among individuals in the population create conditions in which the horizontal cultural spread of the slow reproduction trait is successful, resulting in an overall reduced level of fertility.

**Figure 3.**
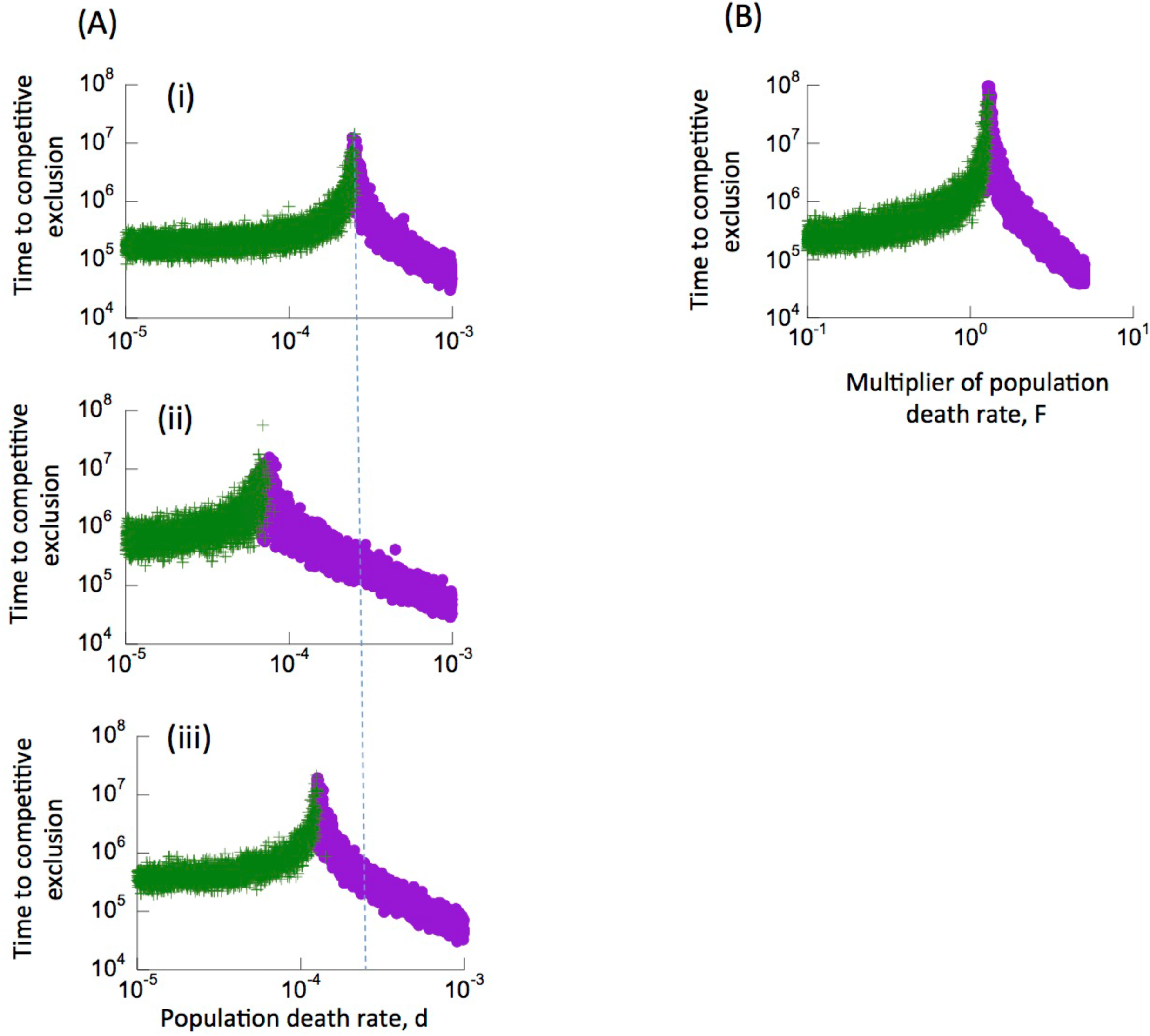
Time to competitive exclusion, as a function of the death rate. (A) Model ABM1. Individual realizations of the computer simulation were run until one of the two populations (fast or slow reproducers) went extinct. This time was recorded with a green dot if the fast-reproducing trait went extinct, and with a purple dot if the slow-reproducing trait went extinct, as a function of the population death rate, D. For low death rates, there are only green dots, corresponding to the slow-reproducing trait persisting and the fast-reproducing trait going extinct relatively fast. For fast death rates, there are only purple dots, corresponding to the opposite outcome. For intermediate death rates, the time until one of the traits goes extinct becomes sharply longer, and either trait can go extinct first. This corresponds to long-term coexistence. For plot (i), parameters were chosen as follows: R_f_=0.005; R_s_=O.8R_f_; P_C_=0.0008; Q=0.93. Plots (ii) and (iii) explore parameter dependence of the phenomenon. (ii) A higher value of Q=0.98 makes it harder for the slow-reproducing trait to invade, hence requiring lower population death rates. (iii) A lower rate of cultural transmission, P_C_=0.0004, makes it harder for the slow-reproducing trait to invade, hence again requiring lower population death rates. (B) Same, but according to ABM2 with age structure. Because each age class is characterized by its own death rate, we multiplied all those death rates by a variable factor F, and plotted the outcome against this parameter. The death rates for the age classes were: D_1_=0.00004; D_2_=0.00007; D_3_=0.00009; D_4_=0.0002. Other parameters are R=0.005; P_C_=0.0008; Q=0.93; A=10,000.

An intuitive explanation is as follows. The death rate determines the rate at which the fast reproducers can outrun the slow reproducers. For large death rates, population density is low and the reproductive potential of individuals is highest. Therefore, fast reproducers can outcompete the slow ones at relatively fast rates, making it difficult for horizontal cultural transmission to reverse this trend. For lower death rates, densities increase, and this slows the rate at which fast reproducers can outrun slow ones. Hence, it becomes easier for horizontal cultural transmission to reverse this process.

Parameters other than the death rate further modulate the outcome of the dynamics, see Figures 3Aii and iii. Cultural transmission of the low fertility trait is promoted by lower values of Q, i.e. by a reduced influence of fast reproducers on choosing the reproduction strategy during the cultural transmission procedure. Increasing the value of Q results in a lower population death rate that is required for cultural transmission to be successful (Figure 3Aii). The relative probability for a cultural transmission event to take place, P_C_, is also an important determinant of outcome. As expected, higher values of P_C_ promote the cultural spread of the fertility-reducing trait. For lower values of P_C_, lower population death rates are needed for cultural transmission to be successful (Figure 3Aiii).

#### (ii) Age structured models: early vs late reproducers

Rather than considering fast versus slow reproducers, we now modify the agent-based model to consider agents who can reproduce either early or late in their lifetimes. This model will be referred to as ABM2. While these two concepts are related, a reduction in fertility due to a later age of first reproduction might be relevant to current times where segments of the population with higher degrees of education and more wealth tend to reproduce at later ages.

In the agent-based model, we consider four age classes. Individuals are born into age-class 1, in which no reproduction is possible. During each time step, all individuals age by one time unit. After A time units, an individual advances to the next age class. Reproduction can occur in age classes 2 and 3 for early reproducers, and only in age class 3 for late reproducers. In either case, reproduction occurs with a probability R. Age class 4 is a post-reproductive phase, during which the only event that can occur is death (the “grandmother effect” has been explored in Section 2.5 of the Supplement; it only influences the main findings in a quantitative way). Death can occur in all age classes, but with increasing probabilities for successive age classes, i.e. with probabilities D_4_>D_3_>D_2_>D_1_.

This model has the same properties as ABM1, see Figure 3B. Some analytical insights for non-spatial, deterministic age-structured models are provided in Supplementary Materials, Section 2.

#### (iii) Continuously increasing population growth, and the simulation of the transition process

Our central result, that a reduction in death rate tends to select for the cultural spread of a fertility-reducing trait, relies on density-dependence in the population dynamics. It is not observed in models assuming unlimited exponential growth, where the rate of cultural transmission alone determines which population outgrows the other. With exponential growth, a reduction in death rate does not slow down the rate at which faster reproducers, by having more offspring, gain advantage over slow reproducers, as was the case with density dependence.

While human population sizes have followed long-term increasing trends, evidence for density-dependent effects and the relevance of local carrying capacities have been found in demographic data from pre-industrial European populations within individual settlements [27]. Continued population growth would then be brought about by an increase in the number of settlements or by regular increases in the carrying capacity, due to advances in society [27,28].

To capture the patterns reported in reference [27], we consider a growing population that is subdivided into neighborhoods or demes (settlements). In each deme, we impose a carrying capacity and describe the local dynamics by ODE model (1). As initial conditions, a single deme is populated with a majority of fast reproducers and a minority of slow reproducers. At the end of each time unit, individuals in each deme have a chance to found a new, empty deme into which a fraction of the current local population moves. The probability of this occurring is proportional to how full the current deme is. This corresponds to an effective increase in population size due to new advances. In addition, the probability to found a new deme is inversely proportional to the number of existing demes. While the demes are not arranged spatially in this model, founding a new deme can be thought of as an increase in the density of the population, which gets more difficult the more demes already exist. Hence, the probability for members of an individual deme to found a new deme is given by 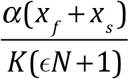, where N is the number of currently populated demes, x_f_ and x_s_ represent local population sizes of fast and slow reproducers, K is the local carrying capacity, and α and ε are constants. When a new deme is founded, a fraction *f* of both fast and slow reproducers moves into the new deme. As more demes become populated, the same algorithm is applied to every deme after each time unit. While in this model, the local dynamics are described by ODEs, it is still a spatial model due to the assumed patch organization, and this approach is consistent with the documented notion of local carrying capacities [27].

In this model, we observe persistence of one trait and exclusion of the other, while the population continues to grow (Figure 4A, B). As before, the fast-reproducing trait persists for high overall death rates (Figure 4A), while the slow-reproducing trait persists for low overall death rates (Figure 4B).

**Figure 4.**
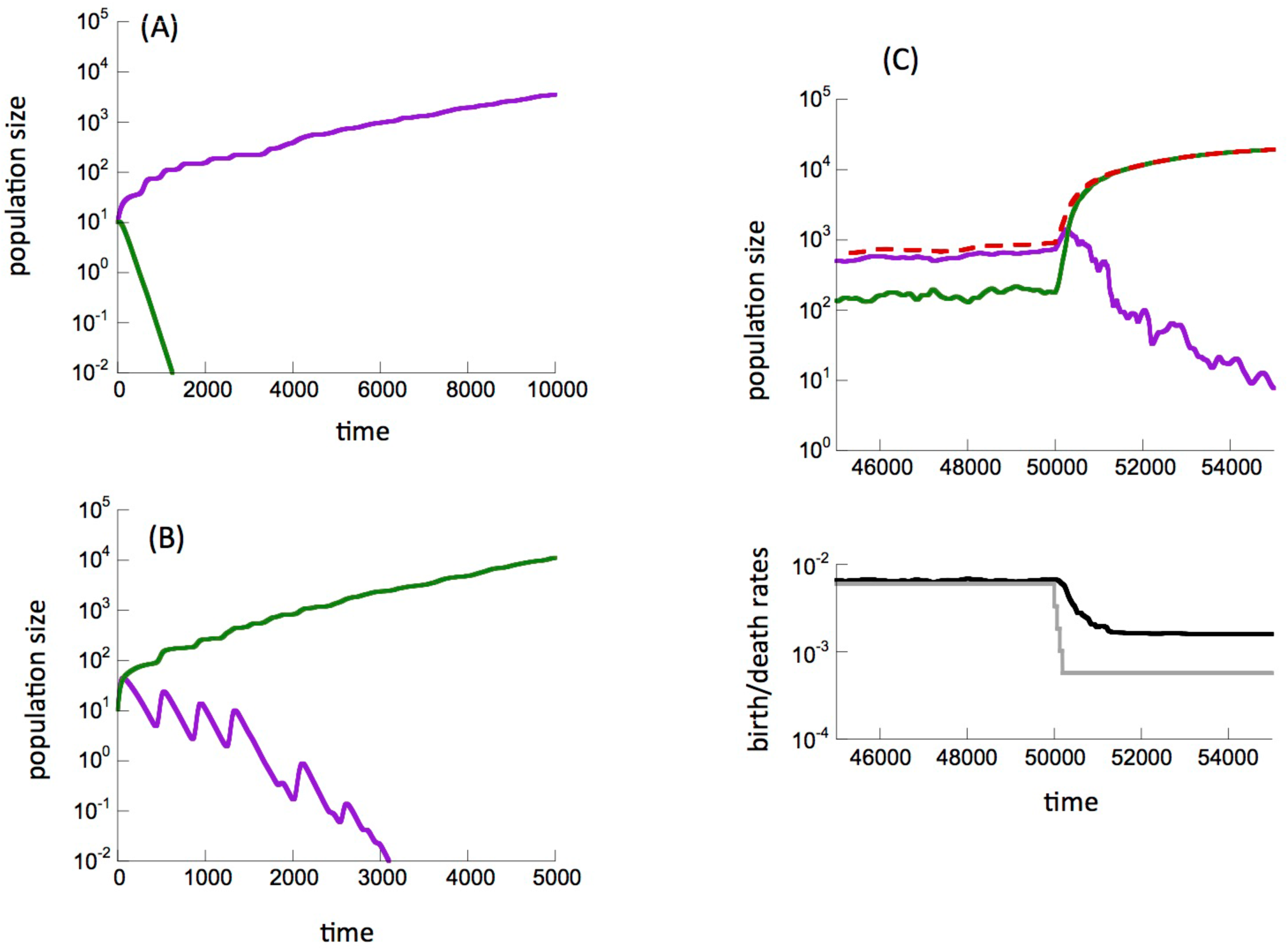
Computer simulations of the deme model, described in the text. (A) The slow-reproducing population (green) goes extinct and the fast-reproducing population (purple) continues to grow. Parameter values were chosen as follows: r_f_=0.08, r_s_=0.064, d=0.05, β=0.01, K=100, α=0.001, ε=0.001. (B) The fast-reproducing trait is going extinct, and the slow-reproducing trait takes over and continues to grow. The same parameter values were used, except d=0.005. (C) Simulation of the demographic transition process. Again, fast- and slow-reproducing traits are shown in purple and green, respectively. The total population size is shown by the dashed red line. The simulation is started with a death rate d=0.006. In this regime, the fast-reproducing trait has an advantage and is dominant. The cultural spread of the low-fertility trait is not successful. At a defined time point, the death rate is reduced 1.8 fold every half generation until it has fallen to a value of d=0.001 (grey line). This creates conditions under which the cultural transmission of the fertility-reducing trait is successful, and the population characterized by a slow reproduction rate spreads. This leads to a decline in the average reproduction rate of the population (black line), which is delayed with respect to the reduction in the death rate. For the parameter regime considered, the average reproduction rate is halved within about 2-3 generations, which corresponds to about 50-100 years (a generation in the model is given by 1/r). The remaining parameters are given as follows. r_f_=0.008, r_s_=0.0016, β=0.2, K=100, α=0.005 ε=0.01.

We further used this model to simulate the demographic transition process (Figure 4C). The simulation was run as before, except that at a defined time point in the simulation, the death rate was continuously and gradually reduced over several time steps towards a lower, new level (Figure 4C, lower panel). This exogenous reduction is shown by the grey line and is assumed to correspond to an improvement in various socio-economic factors that reduce mortality, such as an improvement in disease treatment, sanitary conditions, technological innovations.

In the upper panel, the fast-reproducing population is shown in purple, the slow-reproducing population in green, and the total population size is shown by the red dashed line. Initially, the overall population death rate is relatively high, and the fast-reproducing individuals enjoy a growth advantage. The average reproduction rate is shown by the black line (Figure 4C, lower panel) and is driven by the fast-reproducing population. The overall growth rate of the population is relatively slow at this stage because of the high death rate.

When the death rate is reduced, the fertility-reducing cultural trait can spread successfully and eventually becomes dominant. As the death rate declines, a phase of faster population growth occurs, as observed in data on demographic transitions [22]. Following a time delay after the reduction in the death rate, the average reproduction rate also declines, which is again consistent with data on demographic transitions [22] (Figure 4C, lower panel, black line).

The exact timing of events depends on model parameters. For the purpose of this simulation, we chose parameters such that it takes about 3 generations to reduce the average reproduction rate two-fold. This is an order of magnitude that is similar to events observed in human populations [1] and shows that the cultural transmission dynamics underlying our model can lead to sufficiently rapid changes in fertility. A faster rate of horizontal cultural transmission (higher value of β) can lead to more rapid changes in fertility following the decline in the death rate.

To show that these dynamics are not dependent on this particular model formulation, we performed similar simulations with an age-structured model where continued population growth was allowed through regular increases in the carrying capacity parameter (rather than increasing the number of demes). Similar results were observed and are presented in the Supplementary Materials (Section 2.4).

### 3C. Long term cultural evolution: reproduction strategies as a continuous trait

So far, we considered two distinct populations of slow and fast (early and late) reproducers. To study longer-term evolution, rather than considering two discrete reproductive strategies, it is more realistic to assume the probability of reproduction to be a continuous variable. Because this is most easily implemented in terms of an agent-based model, we will build on the spatial agent-based model of section 3Bii. We again assume that an individual is chosen for a horizontal cultural transmission event with a probability scaled with P_C_. In this model, however, instead of adopting (or rejecting) the reproductive probability of the alternative type, the individual adopts the weighted average of the reproduction probabilities among all neighbors (including its own reproduction probability). As in the above models, we assume that slower reproducers are more influential and contribute more to horizontal cultural transmission than faster reproducers. Due to the continuous nature of the reproduction trait in the current model, this is now implemented during the averaging procedures across the neighborhood: we weigh the reproduction probability by a factor Q<1 if the reproduction probability of a neighbor is faster than that of the individual under consideration.

The outcome observed in this model is straightforward. As initial conditions, the individuals in the system are characterized by different reproduction probabilities. Over time, the reproduction probabilities converge to a spatially uniform value, the level of which depends on the initially assigned probabilities. This eventual uniformity derives from the assumption that an individual adopts the average reproduction probability of the neighborhood during a cultural transmission event.

Next, we introduce mutations of cultural traits that can occur during horizontal transmission. Instead of simply adopting the (weighted) average strategy of the neighborhood, with probability *u* individuals would modify this strategy by increasing or decreasing it (with equal probabilities) by a fraction G. We examined the evolution of the average reproduction probability, R, over time, by running computer simulations. Three types of outcomes were observed (Figure 5).

i. The average probability to reproduce, R, increases steadily towards the maximum possible value (R+D=1), shown by the purple, green, and red lines in Figure 5. (Simulations were stopped when R+D=1).
ii. The average probability to reproduce declines steadily, eventually resulting in population extinction, shown by the dark blue, light blue, and pink lines in Figure 5. Extinction occurs because the reproduction rate evolves to levels that are too low to maintain the population.
iii. The average probability to reproduce converges to an intermediate level, and fluctuates around this level, shown by the yellow and orange lines in Figure 5. This level is independent of the starting value of R (not shown).

**Figure 5.**
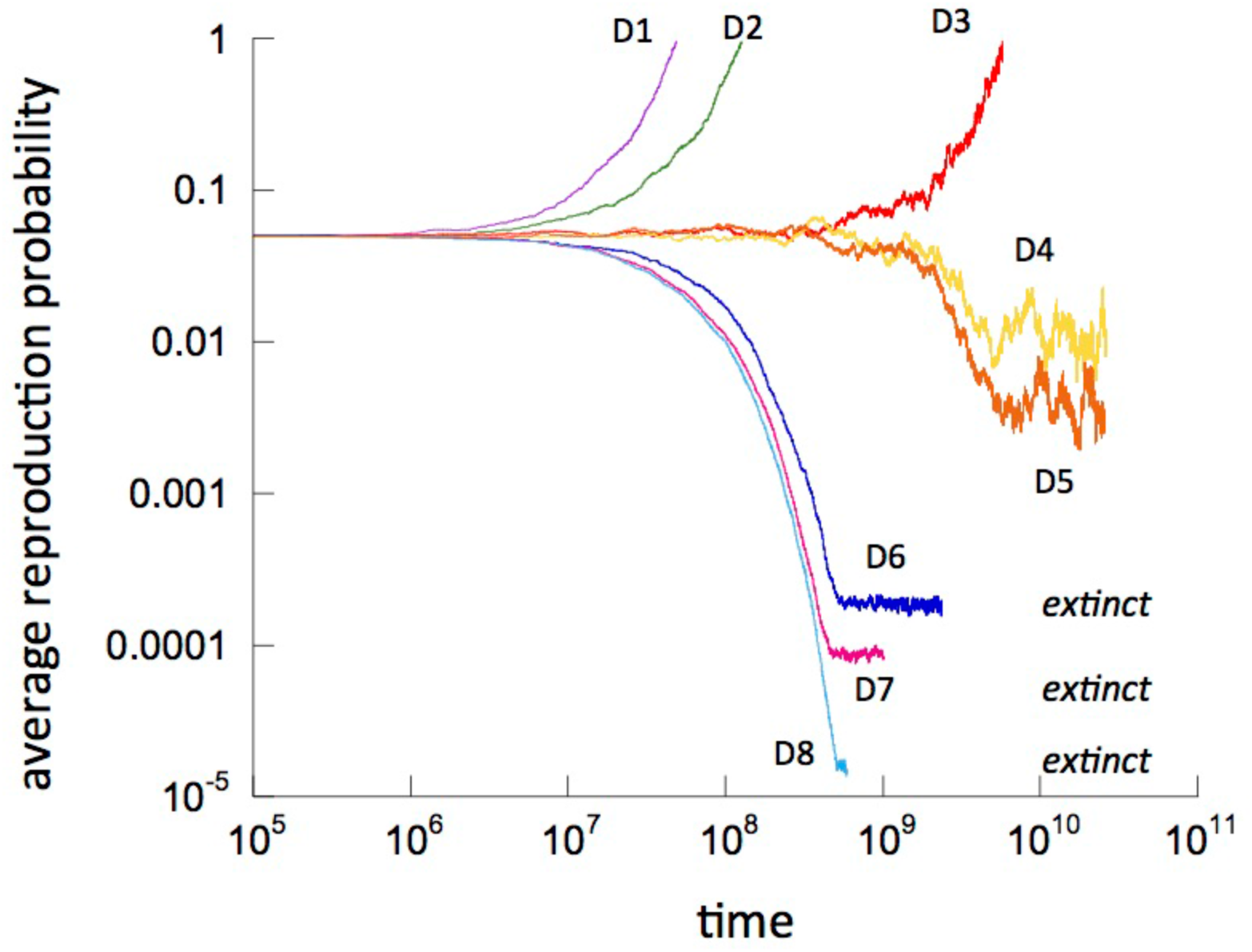
Outcomes of ABM3 with a continuous reproduction strategy and cultural evolution. The average reproduction probability across the whole population is plotted over time. Individual simulation results are shown. Simulations were run for different death rates, decreasing from D1 to D8. For relatively high death rates, the average reproduction probability increases steadily towards maximal levels. For relatively low death rates, the average reproduction probability decreases steadily until population extinction occurs (due to the limited reproduction). For intermediate death rates, the average reproduction probability comes to oscillate around a steady value, which does not depend on initial conditions (not shown). Parameters were chosen as follows. Death rates are given by D1 = 0.002, D2 = 0.001, D3 = 4×10^-4^, D4 = 3.75×10^-4^, D5 = 3.6×10^-4^, D6 = 10^-4^, D7 = 5×10^-5^, D8 = 10^-5^. The reproduction probability of the individuals, R, was allowed to evolve, starting from R=0.05 for all individuals. P_C_=0.0003; Q=0.965. The chance to make a mistake during horizontal cultural transmission (“mutation”) u=0.1. In case of a mistake, the average reproduction rate was changed by G=2%.

As before, the population death probability, D, is a crucial factor (Figure 5). Evolution to maximal reproduction probabilities, R, is seen for relatively large death rates. Evolution towards low values of R and hence population extinction is observed for relatively low death rates. This could be the cultural equivalent to “evolutionary suicide” or “Darwinian extinction” [29]. Evolution towards an intermediate reproduction probability is observed for intermediate death probabilities, D. A higher probability of cultural transmission, P_C_, and a lower weight of faster reproducers during the averaging process, Q, further promote evolution towards declining reproduction rates and population extinction (not shown). Section 3 of the Supplement further explains the existence of an equilibrium state and explores how the mean population reproduction rate depends on parameters.

This model demonstrates that manipulating the death rate changes the long-term cultural evolution of reproductive strategies, and that three different outcomes are possible: the two extremes (maximum reproduction and decline of reproduction rate below replacement level), as well as an evolutionary stable intermediate average reproduction probability. The latter has perhaps been most relevant for human societies, although the trajectories might be moving towards the decline below replacement levels, which is discussed further below. We note that these results were derived from a spatially explicit model. An equivalent non-spatial model is explored in the Supplementary Materials, section 3.1. In the non-spatial model, an evolutionary stable intermediate average reproduction probability is not observed, demonstrating that this outcome depends on the existence of spatially explicit interactions. Finally, the Supplementary Materials (Section 4) further demonstrate that conclusions described here remain robust in a model that assumes sexual reproduction.

## Discussion and Conclusion

We have used a variety of modeling approaches to investigate the basic dynamics by which a fertility-reducing trait can spread via cultural transmission. In contrast to previous modeling approaches, we have allowed for the possibility of exogenous external influences on the population mortality rate. This exogenous parameter can be modulated as a consequence, for example, of technological development in the society. A central result was that lower population death rates select for the cultural spread of the low-fertility trait. This happens because lowering the mortality increases density, which in turn reduces the rate at which the fast reproduction trait gains in abundance relative to the slow reproduction trait. This allows horizontal transmission to tip the balance in favor of slow reproduction. The advantage of the fast reproduction trait is greater when generational turnover is rapid owing to a high death rate. When the death rate declines, there is more opportunity per generation for cultural transmission to operate in favor of the low reproduction trait. We note that the dependence of outcome on population mortality requires the assumption of a cultural transmission bias: individuals with lower reproduction rates need to carry more social weight, an assumption that has also been made in previous modeling work [19]. While it seems reasonable to assume that economically more successful individuals carry more weight in cultural transmission than individuals who are less successful [30,31], the details of this are not well understood [32,33] and require further investigation.

Competition among individuals in the form of density-dependent dynamics was a major driving force underlying the dynamics arising from the model. While in the simpler settings explored here, competition correlated with populations being close to carrying capacity, we showed how a deme model or an age-structured model with increasing carrying capacity can give rise to the same outcomes in populations that continuously grow. Hence, the results described throughout the paper hold for growing populations. We demonstrated that, depending on parameters, the model can reproduce crucial features of the “demographic transition model” [22].

Our study complements previous mathematical work that analyzed the cultural spread of small family sizes in relation to demographic transitions [18–20]. Our models consider a simpler setting involving the basic spread dynamics of the fertility-reducing trait, somewhat similar to infection models. We show that lower death rates promote the cultural spread of the low fertility trait. This result offers a simple possible explanation for the key observation that a reduction in fertility tends to be preceded by a reduction in mortality.

In addition, our model can help interpret demographic data demonstrating that fertility is density dependent [34]. Lowering the death rate in the model leads to an expansion in the slow reproduction trait, even in the context of increased resource availability and continuously growing populations. Data indicate that human fertility as well as family size preference are characterized by density dependence, even during the time frames when demographic transitions occurred. Our model results might offer an explanation for this observation [34].

Also consistent with stylized facts, our models implied that for low population death rates, the average reproduction rate of the population can decline to levels that do not sustain a stable population. In Western European countries, fertility has declined below replacement levels since the 1970s and 1980s [5,6]. Similar tendencies are observed in Japan, South Korea, Taiwan, Singapore, and Hong Kong [35]. In addition, recent surveys [6] have revealed that the mean ideal family size (MIFS) in German speaking countries has fallen below replacement levels, about 1.7 children, among younger people, indicating that this trend might continue in the future. In Taiwan, among women aged 18–24, the MIFS declined from 2.1 in 1993 to 1.8 in 2003, and in Hong Kong, among women aged 18–27, MIFS fell from 1.8 in 1991 to 1.5 in 2011 [36,37].

The models studied here contain a number of assumptions that we consider to be central to exploring the effect of the population death rate on the spread of a culturally transmitted, fertility-reducing trait. Further assumptions and processes could be built into the model, and a detailed exploration of this would be an interesting subject of future research. One such aspect is the grandmother effect [38,39], where individuals in later age classes (grandmothers) promote the survival of individuals in younger age classes. We present a basic exploration of this effect in the Supplement (Section 2.5) and found that this only modulates the parameter thresholds where behavioral changes of the models are observed, but does not qualitatively change outcomes. Another interesting aspect to include might be costs associated with early or late reproduction, which likely also does not lead to a qualitative change of our results. Several additional aspects could be quantified in such more complex models, but this would go beyond the focus of the current manuscript.

While some details of the model processes could be formulated in different ways (see Supplement Section 5), we have considered a range of models with different assumptions. In all models, the death rate of the population was identified as a crucial factor that determined whether the fertility-reducing trait could invade. This could have implications for understanding the forces that contribute to the occurrence of demographic transitions and that drive the decline of fertility below replacement levels in developed countries. It would be interesting for future work to integrate these cultural evolution dynamics with other potential mechanisms that might contribute to the demographic transition process, such as the offspring quality/quantity tradeoff or other economic considerations that might result in human populations having an optimal, target number of offspring.

## Acknowledgements

We would like to thank Simon Levin for useful discussions that helped shape this manuscript.

