## Supplementary Materials for "Evolutionary dynamics of culturally transmitted, fertility-reducing traits"

#### Supplementary information

**4 Sexual reproduction 22**

**5 Model extensions – future work 23**

### 1 In the absence of age structured dynamics

#### 1.1 Alternative ODE models

In the model considered in Section 2.1 of the main text, the conversion process is described by the term

$$\beta x^{(1)} x^{(2)}/K.$$

Alternatively, this term can be formulated as

$$\beta \frac{x^{(1)} x^{(2)}}{x^{(1)} + x^{(2)}}, \tag{1}$$

where the conversion happens proportionally to the current fraction of the individuals of the opposite type. In this case, we have a very similar solution structure. The competitive exclusion solutions are the same as in the previous model, the threshold $d$ values are given by

$$d_1 = \frac{\beta}{1 - r_2/r_1}, \quad d_2 = \frac{\beta}{r_1/r_2 - 1},$$

and the coexistence solution is given by

$$x^{(1)} = \frac{K}{\beta}\left(1 - \frac{\beta}{r_1 - r_2}\right)(d - d_2), \quad x^{(2)} = \frac{K}{\beta}\left(1 - \frac{\beta}{r_1 - r_2}\right)(d_1 - d).$$

In a different modeling approach we assume that conversion happens at the same rate for both strategies, but it is proportional to the weighted fraction of the two strategies in the population. Assuming that strategy 1 is weighed with coefficient $\gamma < 1$, we obtain that the change in numbers for strategy 1 is given by

$$\beta(1 - \gamma) \frac{x^{(1)} x^{(2)}}{\gamma x^{(1)} + x^{(2)}}. \tag{2}$$

In this case, the competitive exclusion solutions are the same as in the previous model, the threshold $d$ values are given by

$$d_1 = \frac{\beta(1-\gamma)}{\gamma(1 - r_2/r_1)}, \quad d_2 = \frac{\beta(1-\gamma)}{r_1/r_2 - 1},$$

and the coexistence solution is given by a somewhat different expression,

$$\begin{aligned} x^{(1)} &= \frac{K}{\beta + d}\left(\frac{\beta}{r_1 - r_2} + \frac{\beta + d}{\gamma r_2 - r_1}\right)\left(\beta + d + \frac{r_1 - \gamma r_2}{\gamma - 1}\right), \\ x^{(2)} &= \frac{K}{\beta + d}\left(\frac{d\gamma}{\gamma - 1} + \frac{\beta(\beta + d - r_1)}{r_2 - r_1} - \frac{(\beta + d)^2 \gamma}{\gamma r_2 - r_1}\right). \end{aligned}$$

### 2 Age structured dynamics

#### 2.1 Model formulation

We will model the competition dynamics of two types that differ by their reproductive strategies. Assume the existence of $N$ discrete age groups for the two types, and denote the abundance of type $s$ in age group $i$ as $x_i^{(s)}$. Reproduction behavior of type $s$ is described by the vector $a_i^{(s)}$, with entries in $[0, 1]$ denoting relative rate of reproduction of this type in age $i$. Individuals of the first type, $s = 1$, correspond to "fast reproducers", and the second type, $s = 2$, to the "slow reproducers" in the previous section. The latter type generally has a tendency to reproduce later than individuals of type 1. In the approach implemented here, type $s$ is characterized by two integers, $i_{start}^{(s)}$ and $i_{end}^{(s)}$, denoting the first and last age groups where reproduction is possible. We have

$$a_i^{(s)} > 0 \ \text{ if } \ i_{start}^{(s)} \leq i \leq i_{end}^{(s)}, \quad a_i^{(s)} = 0 \text{ otherwise},$$

where

$$i_{start}^{(1)} < i_{start}^{(2)}.$$

We can formulate a discrete time dynamical system for these populations as follows:

$$x_1^{(s)}(t+1) = \sum_{j=1}^{N} a_j^{(s)} x_j^{(s)}(t) W, \tag{3}$$

$$\begin{aligned} x_i^{(s)}(t+1) &= w_{i-1}^{(s)} x_{i-1}^{(s)}(t) \left( 1 - \beta_i^{(s)} \frac{\sum_{k=i}^{N} x_k^{(3-s)}(t)}{\sum_{k=i}^{N} (x_k^{(3-s)}(t) + x_k^{(s)}(t))} \right) \\ + &\quad w_{i-1}^{(3-s)} x_{i-1}^{(3-s)}(t) \beta_i^{(3-s)} \frac{\sum_{k=i}^{N} x_k^{(s)}(t)}{\sum_{k=i}^{N} (x_k^{(3-s)}(t) + x_k^{(s)}(t))}, \quad 1 < i \leq N, \end{aligned} \tag{4}$$

where the competition term $W$ can be defined as

$$W = 1 - \frac{\sum_{s=1}^{2} \sum_{k=1}^{N} x_k^{(s)}}{K} \quad \text{or} \tag{5}$$

$$W = \left( 1 + \frac{\sum_{s=1}^{2} \sum_{k=1}^{N} x_k^{(s)}}{K} \right)^{-1}. \tag{6}$$

Equation (3) describes reproduction. Different age groups reproduce with their own rate $a_j^{(s)}$, and the offspring enters age group 1. Equation (4) describes the population moving from age group to age group. Coefficients $w_{i-1}^{(s)}$ describe the probability for an individual of type $s$ to survive until age $i$. The probability of switching type is described by terms including coefficient $\beta$. First we note that expression $3-s$ for $s \in \{1,2\}$ simply returns the type different from type $s$, because $3-s$ gives 2 if $s=1$ and it gives 1 if $s=2$. The probability to switch from type $s$ to type $3-s$ while transitioning to age group $i$ is given by

$$\beta_i^{(s)} \frac{\sum_{k=i}^{N} x_k^{(3-s)}(t)}{\sum_{k=i}^{N}(x_k^{(3-s)}(t) + x_k^{(s)}(t))},$$

and is proportional to the fraction of individuals of age $i$ and older that belong to class $3-s$. With this in mind, we can see that the first term on the right of equation (4) multiplies the probability that an individual does not switch to the other type, and the second term multiplies the probability that switching from $3-s$ to $s$ occurs. System (3-4) assumes no switching at the first stage. To include switching at the first stage, we replace equation (3) with

$$\begin{aligned} x_1^{(s)}(t+1) &= \sum_{j=1}^{N} a_j^{(s)} x_j^{(s)}(t) W \left(1 - \beta_1^{(s)} \frac{\sum_{k=1}^{N} x_k^{(3-s)}(t)}{\sum_{k=1}^{N}(x_k^{(3-s)}(t) + x_k^{(s)}(t))}\right) \\ &+ \sum_{j=1}^{N} a_j^{(3-s)} x_j^{(3-s)}(t) W \beta_1^{(3-s)} \frac{\sum_{k=1}^{N} x_k^{(s)}(t)}{\sum_{k=1}^{N}(x_k^{(3-s)}(t) + x_k^{(s)}(t))}. \end{aligned} \tag{7}$$

#### 2.2 System behavior

System (7, 4) has two exclusion steady states (for $s=1$ and $s=2$), which for competition model (6) are given by

$$x_i^{(s)} = K \prod_{k=1}^{i-1} w_k^{(s)} \frac{r \sum_{m=1}^{N} a_m^{(s)} \prod_{k=1}^{m-1} w_k^{(s)} - 1}{\sum_{m=1}^{N} \prod_{k=1}^{m-1} w_k^{(s)}}, \quad 1 \le i \le N, \tag{8}$$

$$x_i^{(3-s)} = 0, \quad 1 \le i \le N. \tag{9}$$

In figure 1 the behavior of a system with $N=5$ stages is shown. We assumed that for fast reproducers, $i_{start}^{(1)} = 2$, and for slow reproducers, $i^{(2)} = 3$, while $i_{end}^{(s)} = 5$ for both types. For simplicity we assumed that within the reproductive stages, the values $a_i^{(s)}$ were equal to a constant (independent on type and stage). Further, we assumed that the rates $w_i^{(s)}$ were $s$- and $i$-independent, and transfer coefficients $\beta_i^{(s)}$ were $i$-independent (but dependent on $s$).

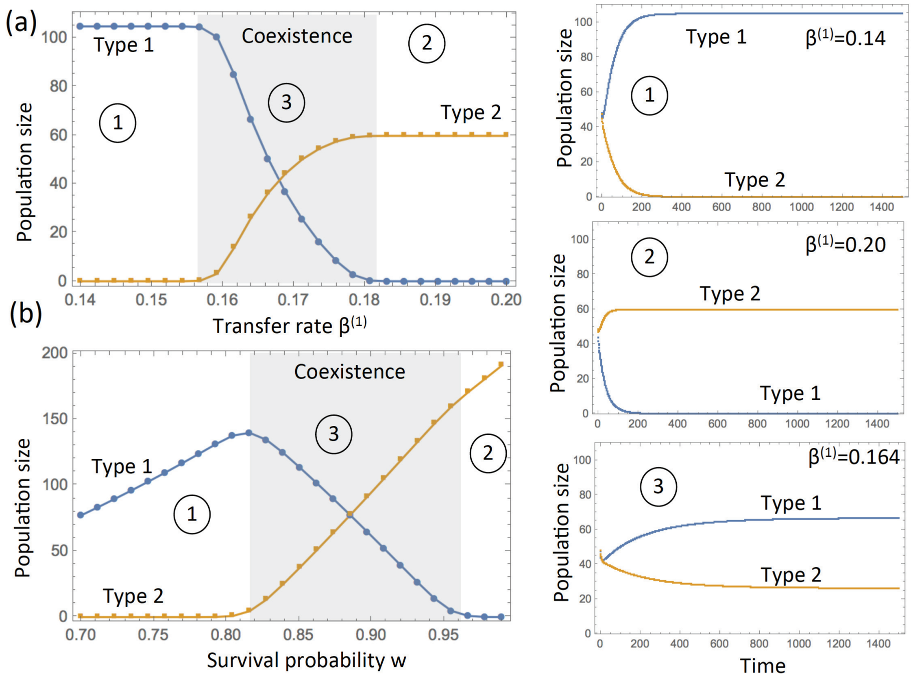

Figure 1: Age structured dynamics according to system (7, 4), numerical simulations. Total populations of individuals of type 1 and type 2 are presented. The steady state values are given on the left as functions of parameter (a) $\beta^{(1)}$ and (b) $w_i^{(s)} = w$ for all $s \in \{1, 2\}, 1 \le i \le N$, the survival probability. Solution types are denoted by a circled number. The parameters are $w = 0.9$ in (a), $\beta^{(1)} = 0.17$ in (b), and $K = 50, \beta_i^{(2)} = 0.1$. The reproductive rate $a_i^{(s)} = 1$ when $2 \le i \le 5$ for $s = 1$ and $3 \le i \le 5$ for $s = 2$. Initially, all populations $x_i^{(s)} = 10$.

In figure 1(a), by fixing all the parameters except for $\beta^{(1)}$, we observed that three different solution types were stable. Solution 1 corresponds to the fast reproducers excluding the slow reproducers and is stable for smaller values transfer away from type 1, $\beta^{(1)}$. Solution 2 corresponds to the slow reproducers excluding the fast reproducers, and corresponds to larger $\beta^{(1)}$. For intermediate values of $\beta^{(1)}$ we observe stable coexistence of both types. Sample time series of the three solution types (corresponding to three different values of $\beta^{(1)}$) are presented on the right on the figure.

Alternatively, if we fix $\beta^{(1)} > \beta^{(2)}$ and vary the survival probability, $w$, the same three solution types are observed, 1(b). In particular, we note that low survival rates (that is, high death rates) lead to the dominance of fast reproducers, and high survival rate (low death rates) to the dominance of slow reproducers.

#### 2.3 A two-age system

The simplest nontrivial system that captures the phenomenon of interest is system (7,4) with $N = 2$. Let us assume that $w_i^{(s)} = w$ for both types (that is, mortality is the same for both types). Further, let

$$i^{(1)}_{start} = 1, \quad i^{(1)}_{end} = 2, \quad i^{(2)}_{start} = 2, \quad i^{(1)}_{end} = 2,$$

in other words, type 1 reproduces both in ages 1 and 2, and type 2 only reproduces in age 2. The trivial solution[1] is unstable if $wa_2^{(2)} > 1$ or $wa_2^{(1)} > 1 - a_1^{(1)}$. The following are some of the non-trivial long-term solutions (compare to the equilibria of section 1):

1. Type 1 (fast reproducers) wins – a competitive exclusion steady state:

$$x_1^{(1)} = \frac{K[r(a_1^{(1)} + wa_2^{(1)}) - 1]}{1 + w}, \quad x_2^{(1)} = \frac{Kw[r(a_1^{(1)} + wa_2^{(1)}) - 1]}{1 + w}, \quad x_1^{(2)} = x_2^{(2)} = 0.$$

2. Type 2 (slow reproducers) wins – a competitive exclusion steady state:

$$x_1^{(1)} = x_2^{(1)} = 0, \quad x_1^{(2)} = \frac{K[ra_2^{(2)}w - 1]}{1 + w}, \quad x_2^{(2)} = \frac{Kw[ra_2^{(2)}w - 1]}{1 + w}.$$

3. A coexistence state.

4. Periodic solutions.

[1] For the analysis of the trivial solution one has to modify the original system by adding a small constant in the denominators of all the equations, otherwise we have a singularity which is meaningless, because the transfer terms multiplying $\beta$ must be zero if the population is zero.

Stability of the two exclusion states can be investigated. For simplicity, let us set all nonzero values of fecundity to a constant, $a_i^{(s)} = a$. Further, we will assume that the coefficient of transfer is independent of the age, and is only defined by the type: $\beta_i^{(s)} = \beta^{(s)}$ for $i = 1, 2,\ s = 1, 2$. Let us analyze stability of solution 1 above (fast reproducers win). Stability of the discrete system requires all the eigenvalues of the Jacobian to satisfy $|\lambda| < 1$. The eigenvalues are given by

$$\lambda_{1,2} = \frac{1 \pm \sqrt{1 + 4rw(1+w)^2}}{2r(1+w)^2}, \tag{10}$$

$$\lambda_{3,4} = \frac{\beta^{(1)}(2+w) \pm \sqrt{w[w(2+\beta^{(1)} - 2\beta^{(2)})^2 + 4(1+\beta^{(1)} - \beta^{(2)})(1-\beta^{(2)})]}}{2(1+w)}. \tag{11}$$

The first two eigenvalues do not depend on the transfer rates and correspond to the stability of the type 1 population in the absence of the other population. We can show that $|\lambda_{1,2}| \le 1$ for all $0 \le w \le 1$ and $r \ge 1$. In particular, $\lambda_1 \ge 0$, we have $\lambda_1 = 1$ when $r = 1, w = 0$, it decays with $r$ and $w$ for $r \le 2$, and for a given $r > 2$, it has a maximum value $(1-w)/2$ when

$$r = \frac{2}{(w-1)^2(w+1)}.$$

Further, $\lambda_2 \in (-1, 0]$ for all values $w \in [0, 1]$ and $r \ge 1$, since $\partial\lambda_2/\partial r > 0$, and for $r = 1$, $\lambda_2 = 1 - \sqrt{1 + 4w(1+w)^2}/(2(1+w)^2) \in [1/8(1-\sqrt{17}), 0]$.

The eigenvalues $\lambda_{3,4}$ describe stability against an invasion of type 2 individuals. The solution can become unstable if $\lambda_3 > 1$. This happens when

$$w > w_1 \equiv \frac{(1-\beta^{(1)})^2}{(\beta^{(2)} - \beta^{(1)})(2 - \beta^{(2)})}.$$

Clearly, if $\beta^{(1)}$ is large (close to 1), the type 1 solution is unstable (because of frequent transfers to type 2). In fact, as long as

$$\beta^{(1)} < \frac{4 - \beta^{(2)} - \sqrt{5(\beta^{(2)})^2 - 16\beta^{(2)} + 12}}{2},$$

the type 1 solution is stable for any values of $w < 1$, because $w_1 > 1$. If however the inequality above is revered (that is, the transfer rate is larger than a threshold for type 1), the solution becomes unstable for sufficiently large values of $w$.

Intuitively, success of each of the types depends on their net fecundity and their propensity to stay (and not transfer to the opposite type). Clearly, the fecundity of type 1 is larger than that of type 2. But this can be offset by a larger probability of transfer (if we assume that $\beta^{(1)}$ is larger than $\beta^{(2)}$ by a sufficient margin). Small death rates (and therefore large values of $w$) work against type 1 individuals and benefit type 2 individuals. If $w$ is large, more individuals survive to later stages, resulting in a larger influx of individuals transferring from type 1 to type 2: they simply have a longer time to stay alive and decide to switch. Thus, living longer increases success of type 2, such that after a threshold of $w$, type 2 becomes stronger and drives type 1 extinct.

Investigating the stability of type 2 equilibrium, we discover that it is unstable (in this simple 2-age model) for all values of $w$ except for $w = 1$, where it is neutral. Note that for systems with more age stages, this is not the case, and we have a stable type 2 equilibrium (see the previous section). For the 2-age system, for values $w < 1$, but close to 1, instead of equilibrium 1, we observe a stable cycle which contains only type 2 individuals.

#### 2.4 Simulating demographic transition

In this section we present an example of an age structured model where a behavior resembling demographic transition can be observed. We use the following formulation:

$$x_1^{(s)}(t+1) = \sum_{j=1}^{N} a_j x_j^{(s)}(t), \tag{12}$$

$$\begin{aligned} x_i^{(s)}(t+1) &= W_i \left( w x_{i-1}^{(s)}(t) \left( 1 - \beta^{(s)} \frac{\sum_{k=i}^{N} x_k^{(3-s)}(t)}{\sum_{k=i}^{N} (x_k^{(3-s)}(t) + x_k^{(s)}(t))} \right) \right. \\ &+ \left. w x_{i-1}^{(3-s)}(t) \beta^{(3-s)} \frac{\sum_{k=i}^{N} x_k^{(s)}(t)}{\sum_{k=i}^{N} (x_k^{(3-s)}(t) + x_k^{(s)}(t))} \right), \quad 1 < j \leq N, \end{aligned} \tag{13}$$

where we defined

$$W_i = \begin{cases} \left(1 + \frac{\sum_{s=1}^{2} \sum_{k=1}^{N} x_k^{(s)}}{K}\right)^{-1}, & i = 1, \\ 1, & i > 1. \end{cases} \tag{14}$$

In this description, the competition term, $W_i$, is interpreted as infant (or early childhood) mortality, and therefore appears as a multiplier in front of the right hand side of the equation for age group 1, modifying the probability of survival until this stage.

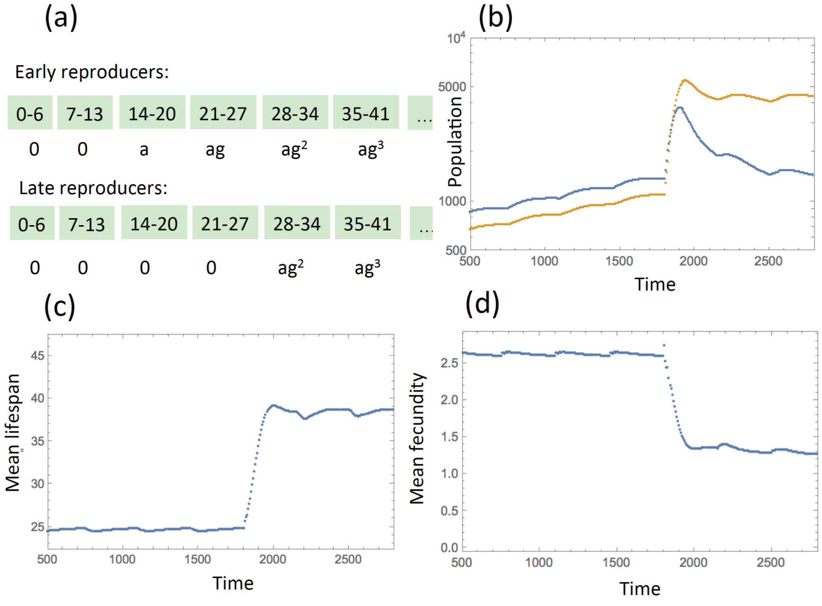

Figure 2: Simulations of an age-structured, deterministic model. (a) Age stages with early and later reproducers' age-specific fecundity are specified. (b) Simulation results for the total population sizes of early (blue) and late (yellow) reproducers, as functions of time, in a simulation with step-wise increasing carrying capacity ($K_T = 1.15\, K_{T-1}$, where $T$ counts periods of 50 age-stages, which is 350 years). The survival probability $w$ is increased exogenously, in a step-like manner at time 1800 from $w = 0.8$ to $w = 0.95$, marking the beginning of a change similar to demographic transition. (c) The mean life-span of the population corresponding to the same simulation is shown as a function of time. (d) The mean number of offspring per individual is shown as a function of time. The rest of the parameters are: $N = 13$ age stages, $a = 2$, $g = 0.5$, $\beta^{(1)} = 0.4$, $\beta^{(2)} = 0.37$.

The probability to survive to the next stage is assumed stage- and type-independent, $w_i^{(s)} = w$ for $s = 1, 2$ and all $i$. Further, the rate of conversion is stage-independent ($\beta_i^{(s)} = \beta^{(s)}$ for all $i$) and parameter $a_i^{(s)}$ related to fertility is type-independent ($a_i^{(s)} = a_i$ for $s = 1, 2$).

Figure 2 presents numerical simulations of model (12-13). For these simulations, we considered 13 age-stages, which represent age groups $0 - 6, 7 - 13, 14 - 20$, etc. There are two types of individuals: early reproducers reproduce in stages 3-8, and later reproducers only reproduce in stages 5-8. Each stage is characterized by the mean age-specific fecundity, which decays exponentially with age and is given by parameter $a$ at stage 3, and by $ag^{k-3}$ at stage $3 < k \leq 8$, where $0 < g < 1$, see panel (a) of figure 2.

In the simulation, we assumed that the carrying capacity, $K$, that defines the maximum population size increases in a step-wise manner. This process is an idealization meant to simulate human expansion. In a space-free model it can correspond both to an increase in density and an outward expansion. In a spatial, agent-based model, a similar effect could be achieved by refining the grid, making it more and more dense. The reason to simulate expansion by increasing $K$ instead of using a model with exponential (uninhibited, non-density dependent) growth is the notion of competition for resources and crowdedness, which are assumed to be important factors in human population dynamics. The population continues to grow through expansion, innovation, and making more resources available, but at the same time the effects of increasing density and frequent resource shortages are felt through density-dependent factors in the equations. In the current model, the density-dependent factors are presented as term $W_1$ entering as infant and childhood mortality factor.

In order to simulate an improvement in mortality, we assumed that the survival probability, $w$, increases in a step-like manner at year 1800 in the simulated system. Figure 2(b,c,d) shows numerical simulations of system (12-13). Panel (b) plots the total population sizes of early (blue) and late (yellow) reproducers as functions of time. Before the transition, the population contains a majority of early reproducers; the mean lifespan is about 25 years (panel (c)) and the mean total fecundity is above 2.5 children per individual (which in a sexually reproducing population would translate into over 5 children per woman). After the transition, the population experiences an increased growth followed by a slow-down (panel (b)). The population now consists predominantly of slow reproducers, the mean life-span increases to over 40 years, and the mean number of children drops to about 1 (equivalent to 2 children per woman). The transition happens on a relatively short time-scale equivalent to under 5 generations.

While the time-scale of the transition, mean longevity and fecundity, as well as growth rate of the population are defined by model parameters, the above simulations demonstrate that an effect similar to demographic transition can be observed in the model, and that parameters can be found such that some of the observables are not far from their realistic ranges.

#### 2.5 Including the "grandmother hypothesis"

To include the so called "grandmother effect", we note that help of a grandparent can increase the chances of a child's survival. To incorporate this we will use system (3-4) as a basic model. For simplicity, we will keep the description asexual. As an individual ages, it passes through stages, and the probability to survive from age $i-1$ to age $i$ is given by $w_{i-1}$ (here we for simplicity assume no explicit dependence of mortality on type $s$). Note that in many contexts,

$$w_1 < w_2.$$

Here we assume that the presence of the grandmother may increase the probability of survival during the earliest stage. Let us denote the probability to have a grandmother by $P_{grand}$. Then we can set the probability of survival to age group 2 to be

$$w_1 + S_{grand}P_{grand}(w_2 - w_1), \tag{15}$$

where $S_{grand}$ is a tunable parameter that sets the strength of the "grandmother effect". If this effect is nonexistent ($S_{grand}P_{grand} = 0$), then mortality of age group 1 is simply $w_1$. If $P_{grand}S_{grand} = 1$, then the probability to survive the first age class is as high as that for the next age class ($w_2$).

To calculate the probability to having a grandmother, we note that for a given newborn, this depends on the age of its parent. Having a younger parent increases the probability that the grandparent is alive. To incorporate this effect, we must use a more detailed description compared to system (3-4), and as the basic variable use

$$y_{ij}^{(s)}(t),$$

which is the number of individuals of type $s$ of age $i$ born to a parent of age $j$, at time $t$. These are related to the old variables $x_i^{(s)}(t)$ as

$$x_i^{(s)} = \sum_{j=i_{start}}^{i_{end}} y_{ij}^{(s)}(t),$$

where we denoted

$$i_{start} = \min\{i^{(1)}_{start}, i^{(2)}_{start}\}, \quad i_{end} = \max\{i^{(1)}_{end}, i^{(2)}_{end}\}.$$

The changes in each type are described by the following equations:

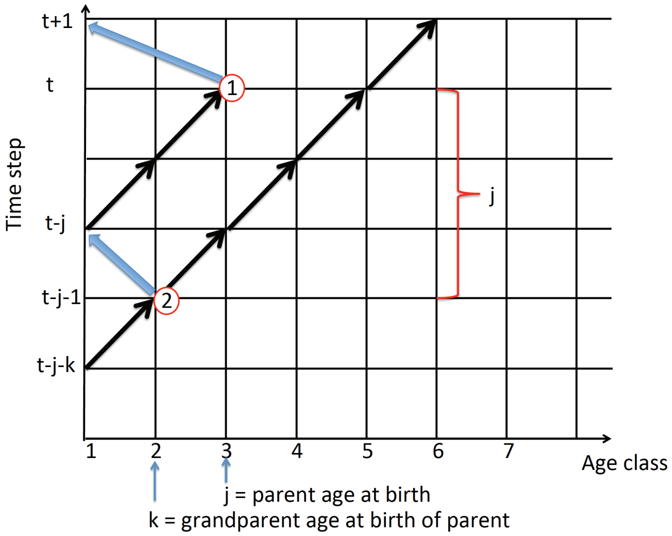

Figure 3: A schematic illustrating the grandmother effect. The two axes are the age class and time. The label "1" represents the birth of an individual, such that at time $t+1$ the newborn enters age-class 1. The age of the parent is $j = 3$ for this example. The black arrows pointing to "1" trace the growing up of the parent. The label "2" marks the birth of the parent to the grandparent of age $k = 2$. The black arrows pointing to the right and upward from "2" represent the aging of the grandparent.

$$\begin{aligned} y^{(s)}_{1j}(t+1) &= a^{(s)}_j x^{(s)}_j(t) W, && (16)\\ y^{(s)}_{ij}(t+1) &= w_{i-1,j} y^{(s)}_{i-1,j}(t)\left(1 - \beta^{(s)} \frac{\sum_{k=i}^N x^{(3-s)}_k(t)}{\sum_{k=j}^N (x^{(3-s)}_k(t) + x^{(s)}_k(t))}\right) \\ &+ w_{i-1,j} y^{(3-s)}_{i-1,j}(t) \beta^{(3-s)}_j \frac{\sum_{k=i}^N x^{(s)}_k(t)}{\sum_{k=i}^N (x^{(3-s)}_k(t) + x^{(s)}_k(t))}, && (17)\\ & 1 < i \le N, \quad i_{start} \le j \le i_{end}, \quad s = 1, 2. \end{aligned}$$

Let us set all the values $w_{ij} = w_2$ for all $i \geq 2$, and assume that the information about the grandmother effect is included in the mortality rate of the youngest age class, $w_{1j}$. We have (as in formula (15)),

$$w_{1j} = w_1 + S_{grand} P^j_{grand}(w_2 - w_1),$$

and the probability of having a grandmother depends on the age of the individual's parent, $j$. To calculate this we use the diagram of figure 3. If the parent's age at time $t$ is $j$ and the parent was born to the grandparent of age $k$ (at time $t - j - 1$), then at time $t + 1$, the age of the grandparent is given by $k + j + 1$. The probability that the grandparent survives to time $t + 1$ is given by the product of probabilities to survive from age $k$ to age $k + j + 1$,

$$\prod_{m=k}^{k+j} w_m,$$

where we assume that $w_m = 0$ for $m \geq N$. The probability of having a grandparent is then given by

$$P^{j,s}_{grand} = \frac{\sum_{k=1}^{N} a_k^{(s)} x_k(t - j - 1) \prod_{m=k}^{k+j} w_m}{\sum_{k=1}^{N} a_k^{(s)} x_k(t - j - 1)},$$

which is the probability that at the moment of the parent's birth the grandparent was young enough to survive to the birth of grandchild (time $t + 1$). Note that this expression makes system (16-17) non-local, that is, the equations now depends on the variable's value in the past (time $t - j - 1$).

In this version of the model, any grandparent that survives can contribute to the increased survivability of the newborn ("strong grandmother effect"). In a different version, we can assume that only grandparents that can no longer reproduce themselves participate in the care for their grandchildren ("weak grandmother effect"). In this case, we require that the age of the grandparent at time $t$, $k + j > i^{(s)}_{end}$:

$$P^{j,s}_{grand} = \frac{\sum_{k=i^{(s)}_{end}-j+1}^{N} a_k^{(s)} x_k(t - j - 1) \prod_{m=k}^{k+j} w_m}{\sum_{k=1}^{N} a_k^{(s)} x_k(t - j - 1)}.$$

Figure 4 shows how model behavior changes as we include strong or weak grandmother effect. In this example, there are $N = 7$ stages, and early reproducers (type 1) start reproducing in stage 2, while type 2 start at stage 3. Individuals do not reproduce in stages 6 and 7. In the absence of grandmother effect, for low values of $w$, type 1 individuals dominate (and exclude type 2); for high values of $w$ the situation is reversed, and for intermediate $w$ we have coexistence of both types. Adding the grandmother effect does not change this picture qualitatively, but gives type 1 individuals a larger advantage, such that the transition to the dominance of type 2 happens for higher values of $w$.

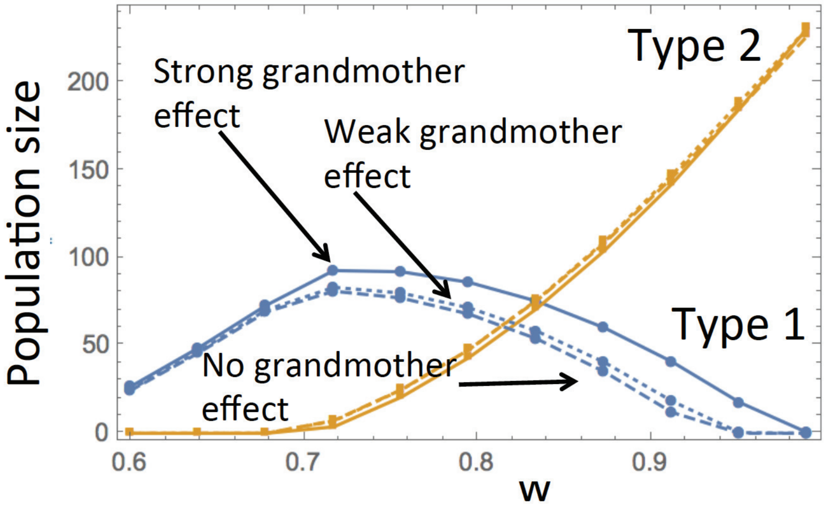

Figure 4: Age structured dynamics according to system (16-17), numerical simulations (similar to figure 1(b)), where the grandmother effect was included. Total populations of individuals of type 1 and type 2 are presented. Dashed lines correspond to no grandmother effect, dotted lines to the weak grandmother effect, and solid lines to strong grandmother effect. The parameters are: $N = 7$ age stages, $i^{(1)}_{start} = 2, i^{(2)}_{start} = 3, i^{(1)}_{end} = i^{(2)}_{end} = 5; w_1 = 0.9w, \beta^{(1)} = 0.2, \beta^{(2)} = 0.1, K = 140, S_{grand} = 1, a^{(s)}_i = 1$ during reproductive stages.

### 3 Birth-death, imitation, space, and mutation dynamics

#### 3.1 Differences between spatial and non-spatial models.

Here we illustrate that spatial and non-spatial modeling approaches yield qualitatively different results in the context of long-term evolutionary dynamics of reproductive strategies.

In this paper, we consider two broad types of models. The first type of model analyzes the competition between slow versus fast reproducers. As mentioned, spatial interactions do not change the nature of the results. The second type of model considers the reproduction rate as a continuous trait, and analyzes its long-term evolution (section 3C of the main text). Here, spatial restrictions do alter the outcome. We ask to what evolutionary stable reproduction rate the system evolves. In the spatial model (figure 5, main text), we observe 3 different evolutionary trajectories: (i) the average reproduction rate evolves to maximal values, i.e. fastest reproduction. (ii) the reproduction rate evolves to below replacement levels. (iii) the reproduction rate evolves towards an evolutionary stable intermediate state (see the curves corresponding to D4 and D5). In contrast, in an equivalent agent-based model without spatial restrictions, see figure 5 of this Supplement, outcome (iii) is not observed. In other words, the non-spatial system cannot evolve to an evolutionary stable intermediate reproduction rate.

#### 3.2 Simplified model formulation and numerical results

Envisage the following process. In a 1D spatial system of a constant size, $N$, each individual, $i$, is characterize by a reproduction rate, $l_i$. During each time unit, $N$ updates are performed, each consisting of two parts, a death-birth (DB) update and a cultural transmission (CT) update. Each update proceeds as follows:

- A DB update: An individual is chosen, randomly and fairly, to be removed (say, this is the individual at location $i_1$). Then it is replaced by the progeny of one of its two neighbors: the individual at location $i_1+1$ reproduces with probability $l_{i_1+1}/(l_{i_1+1}+l_{i_1-1})$, and the individual

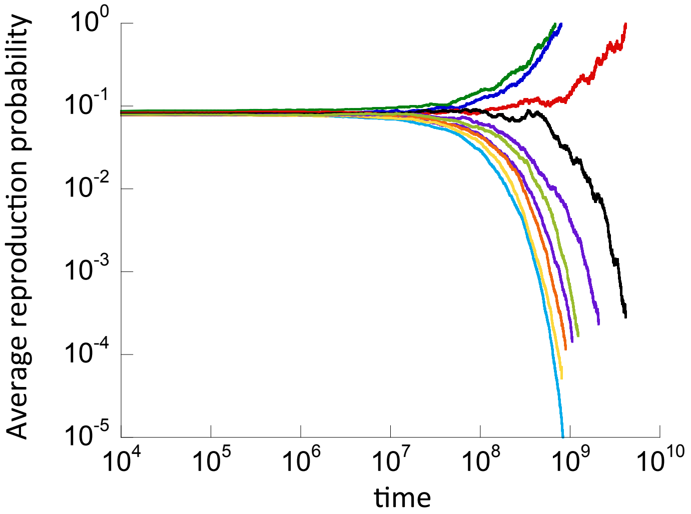

Figure 5: Outcomes of the non-spatial variant of ABM3 with a continuous reproduction strategy and cultural evolution (compare to figure 5 of the main text). The death rates from the top to the bottom curve are: $D1 = 3.75 \times 10^{-4}$, $D2 = 3.6 \times 10^{-4}$, $D3 = 3 \times 10^{-4}$, $D4 = 2.5 \times 10^{-4}$, $D5 = 2 \times 10^{-4}$, $D6 = 1.5 \times 10^{-4}$, $D7 = 1.25 \times 10^{-4}$, $D8 = 1 \times 10^{-4}$, $D9 = 5 \times 10^{-5}$, $D10 = 1 \times 10^{-5}$.

at location $i_1 - 1$ reproduces with probability $l_{i_1-1}/(l_{i_1+1} + l_{i_1-1})$. The offspring inherits the reproduction rate of the parent.

- A CT update: this event happens with probability $\beta$, which sets the relative time scale of the two types of updates. Pick an individual, randomly and fairly, to perform an imitation update (say this is the individual at location $i_2$). This individual will change its reproduction rate from $l_2$ to

$$\tilde{l} = \frac{\sum_{j=i_2-1}^{i_2+1} \alpha_{i_2,j} l_j}{\sum_{j=i_2-1}^{i_2+1} \alpha_{i_2,j}},$$

where

$$\alpha_{i,j} = \begin{cases} 1, & l_j \leq l_i, \\ s, & l_j > l_i, \end{cases}$$

and $0 < s < 1$ is a constant that indicates by how much the strategy of fast reproducers is discounted. In other words, a weighted average of all the strategies around the focal individual at $i_2$ is formed, such that the strategy of those who reproduce faster than the focal individual is discounted with coefficient $s$. The focal individual adopts the resulting strategy with probability $1-u$. With probability $u$, strategy $\tilde{l}$ is increased or decreased (with equal likelihood) by an amount $\Delta l$ (unless $l < \Delta l$, in which case it can no longer decrease). This process is equivalent to mutations, whereby the phenotype is modified with a certain probability to give rise to variation.

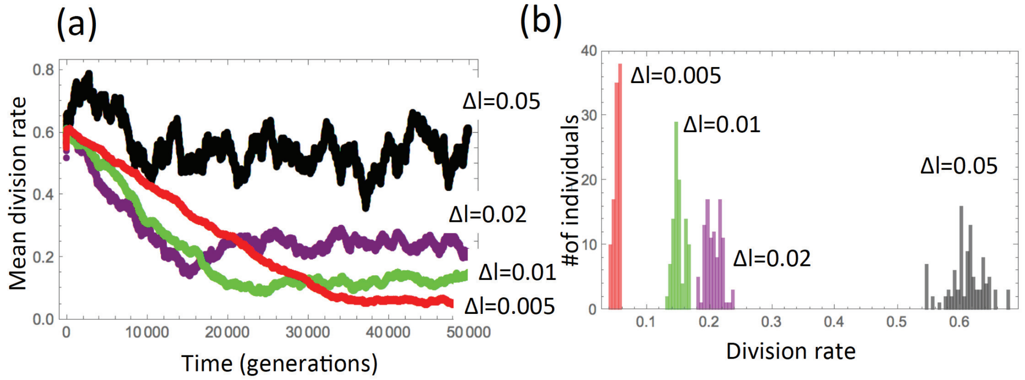

Figure 6: The dynamics of a 1D simulation with mutations. (a) The time-series of the population mean reproduction rate, for 4 different values of $\Delta l$. (b) Numerically obtained histograms of the population's reproduction rates, taken at generation 50,000, for the same 4 values of $\Delta l$. The rest of the parameter are: $N = 100$, $u = 0.04$, $\beta = 1$, $s = 0.9$.

We would like to characterize the equilibrium of this system. First we note that in the absence of mutations ($u = 0$), the state with $l_i = l$ for all $i$ is a equilibrium for any value of $l$. As a result, the system will converge to one of these neutral equilibria, depending, for example, on the initial condition.

The dynamics change drastically in the presence of mutations, $u > 0$. Now, uniform states are no longer equilibrium states, and the equilibrium reproduction rates will be distributed around some mean value, $\bar{l}$, with the variance that increases with $u$ and $\Delta l$. In figure 6(a) we present the time series of the population mean reproduction rates, for 4 different values of $\Delta l$, the increment of the reproduction rate. We can see that the population settles to a stochastic equilibrium, where the mean population mean reproduction rate increases with $\Delta l$, and convergence time decreases with $\Delta l$. Figure 6(b)

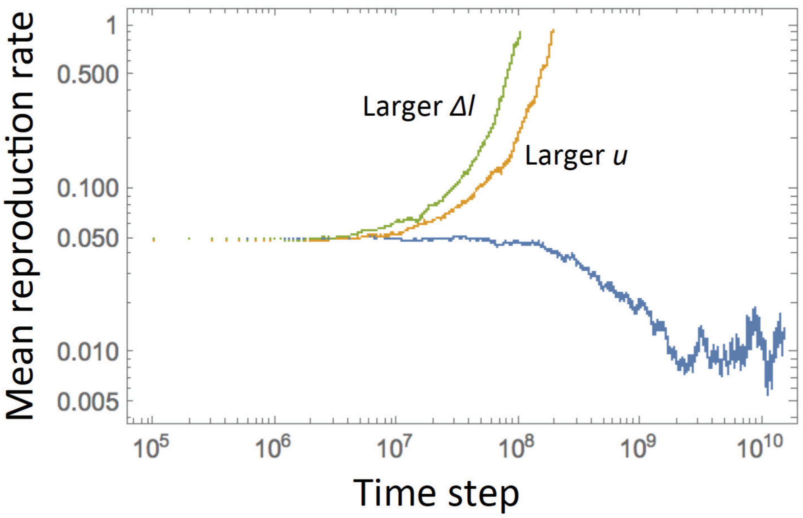

Figure 7: The dynamics of a 2D simulation with mutations. The population mean reproduction rate is plotted as a function of time, for 3 simulations. The blue line represents a base-line simulation with parameters $u = 0.1, \Delta l/l = 0.02$, the orange line a simulation with an increased mutation rate, $u = 0.3$, and the green line a simulation with an increased $\Delta l/l = 0.04$. The rest of the parameters are as in figure 5 of the main text, with the death rate $3.75 \times 10^{-4}$.

shows numerically obtained histograms of reproduction rates of populations at equilibrium, for the same four values of $\Delta l$. We can see that the standard deviation increases with $\Delta l$. Similar trends are observed when we vary the mutation rate, $u$ (not shown). 2D simulations that show the same trends are shown in figure 7.

#### 3.3 Analytical considerations

To find the mean equilibrium value of the reproduction rates, we use the following argument. Suppose that the equilibrium distribution[2] of the reproduction rates is given by $\{f_k\}$, such that the probability for an individual to

[2] A similar argument for continuous distributions can be developed.

have reproduction rate $L_k$ is given by $f_k$, with

$$\sum_k f_k L_k = \bar{l}.$$

Under a BD event, suppose an individual at position $i_1$ with reproduction rate $L_1$ is picked for replacement, and suppose further than its two neighbors have reproduction rates $L_2$ and $L_3$. Then the expected increment in the reproduction rate of the focal individual is given by

$$-L_1 + L_2\frac{L_2}{L_2+L_3} + L_3\frac{L_3}{L_2+L_3}.$$

Averaging over all the possible reproduction rates, we obtain the expected increment in reproduction rate from a DB update:

$$\Delta L_{DB} = \sum_i \sum_j \sum_k \left(-l_i + \frac{l_j^2}{l_j+l_k} + \frac{l_k^2}{l_j+l_k}\right) f_i f_j f_k. \tag{18}$$

Similarly, we can calculate the expected increment in the reproduction rate resulting from a cultural transmission event:

$$\Delta L_{CT} = \sum_i \sum_j \sum_k \left(-l_i + \frac{l_i + \alpha_{ij} l_j + \alpha_{ik} l_k}{1+\alpha_{ij}+\alpha_{ik}}\right) f_i f_j f_k. \tag{19}$$

The equation

$$\Delta L_{DB} = -\beta \Delta L_{CT} \tag{20}$$

characterizes the equilibrium. Note that the right hand side of this equation is positive, because the mean increment resulting from CT updates is negative, due to a diminished weight of high reproduction rates in the weighted averages. The left hand side is also positive, because DB updates tend to increase the reproduction rates due to competition among individuals.

Let us assume that the width of the distribution of the equilibrium reproduction rates is defined by the mutation rate (and the increment $\Delta l$), and keep it fixed, while varying the mean $\bar{l}$. Note that in equation (19), the expression in the parentheses can be rewritten as

$$\frac{\alpha_{ij}(l_j - l_i) + \alpha_{ik}(l_k - l_j)}{1+\alpha_{ij}+\alpha_{ik}}.$$

For each location $i$, let us present $L_i = \bar{l} + \epsilon m_i$, where all $m_i$ are IID with a zero mean and a variance that we denote by $(\sigma/\epsilon)^2$. We can see that $\bar{l}$ cancels from the above expression, and its statistics will only depend on the distribution width. In other words, the mean decrement received by the population reproduction rate as a result of a CT update is defined by the difference between the focal reproduction rate and a weighted average of its neighboring reproduction rates, and does not depend of the absolute value of the rates.

On the contrary, the DB increment defined by equation (18) depends on the magnitude of $\bar{l}$. Intuitively, neighbors compete for filling the empty spot, and the amount of advantage experienced by a neighbor with a higher reproduction rate is proportional to the relative, and not absolute, difference in the rates. Therefore, the increment scales with the relative amount of spread in reproduction rates, and is thus inversely proportional to $\bar{l}$. Again, for each location $i$, we present $l_i = \bar{l} + \epsilon m_i$, where all $m_i$ are IID with a zero mean and variance $(\sigma/\epsilon)^2$. Then, expanding the expression in parentheses in (18) in terms of $\epsilon$ we obtain

$$\left(\frac{m_j + m_k}{2} - m_i\right)\epsilon - \frac{\epsilon}{2}\frac{(m_j - m_k)^2}{m_j + m_k}\sum_{n=1}^{\infty}\left(-\frac{(m_j + m_k)\epsilon}{2\bar{l}}\right)^n.$$

The first term averages to zero, and the second term is given by

$$\frac{\epsilon^2}{4\bar{l}}(m_j - m_k)^2,$$

which upon averaging yields

$$\frac{\sigma^2}{2\bar{l}},$$

a quantity inversely proportional to the mean reproduction rate of the population. We further see that it depends on the square of $\sigma$ in the lowest order.

From the above analysis it follows that the left hand side of equation (20) is a decaying function of $\bar{l}$ which tends to zero as $\bar{l} \to \infty$, and the right hand side of equation (20) is $\bar{l}$-independent. There will be a unique intersection of the two curves as long as $\beta$ is chosen to be sufficiently low. This intersection defines the equilibrium value of the population mean reproductive rate.

We further note that the quantities $\Delta_{DB}$ and $-\Delta_{CT}$ both grow with the distribution width of the reproduction rates, but while $-\Delta_{CT}$ is linear in $\sigma$,

$\Delta_{DB}$ is quadratic in this quantity, and thus grows faster as we increase the width of the distribution of $l$. Therefore, as $u$ increases and the distribution width increases, the left hand side of equation (20) grows faster than the right hand side, resulting in an increase in the solution, $\bar{l}$.

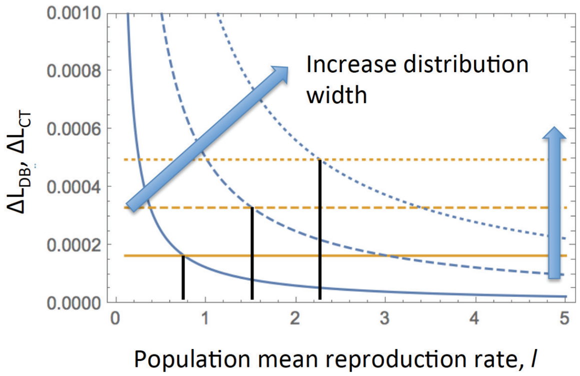

Figure 8: Finding the equilibrium reproduction rate by solving equation (20), illustrated with example (21-22). The left hand side of equation (20), $\Delta L_{DB}$, is shown as blue lines and the right hand side, $-\beta \Delta L_{CT}$, with yellow lines, as functions of $\bar{l}$. Solid, dashed, and dotted lines correspond to three different values of $\Delta l$: $0.05, 0.10, 0.15$. The rest of the parameters are: $s = 0.9, \mu = 0.1, \beta = 1$.

This is illustrated in an example where we assumed that the division rates are distributed according to the following three-valued distribution with mean $\bar{l}$ and variance $(\Delta l)^2 \mu$:

| $i \rightarrow$ | 1 | 2 | 3 |
|---|---|---|---|
| $L_i$ | $\bar{l} - \Delta l$ | $\bar{l}$ | $\bar{l} + \Delta l$ |
| $f_i$ | $\mu/2$ | $1 - \mu$ | $\mu/2$ |

The expressions for $\Delta L_{DB}$ and $\Delta L_{CD}$ can be obtained explicitly,

$$\Delta L_{DB} = \frac{(\Delta l)^2 \mu}{2\bar{l}} \frac{(\Delta l)^2 \mu - 4\bar{l}^2}{(\Delta l)^2 - 4\bar{l}^2}, \qquad (21)$$

$$\Delta L_{CT} = \frac{\Delta l \mu (1-s)}{6(2+s)(1+2s)} \left((6-\mu)\mu s - 10s + \mu(3+\mu) - 8\right). \qquad (22)$$

In figure 8, both sides of equation (20) are plotted as functions of $\bar{l}$, and their intersections are marked with vertical lines, for three values of $\Delta l$, which represent an increase in the distribution width. We can see that the corresponding solutions $\bar{l}$ become larger for larger distribution widths.

### 4 Sexual reproduction

For greater realism, model ABM4 introduces sexual reproduction. ABM4 is based on model ABM3, in that it assumes the probability of reproduction to be a continuous trait, and also that for a cultural transmission updates, a given individual adopts the weighted average reproduction probability of the neighborhood with the possibility of "mutations" as defined in the main text for ABM3. Sexual reproduction is incorporated in the following way. Two genders are distinguished, gender 1 and gender 2. Before reproduction can occur, two individuals of opposing gender have to form an exclusive connection, thus assuming monogamy. The following events can occur if an individual is chosen for a reproductive update. If the individual does not have a partner, a connection can be formed with a probability $M$ if an individual of the opposite gender without a partner is present among the eight nearest neighbors. The partner is randomly chosen from the neighborhood. If the individual does have a partner, reproduction happens with a probability $R_{av}$, which represents the average reproduction probabilities of the two parents. For simplicity, it is assumed that once formed, a partnership cannot break, corresponding to life-long monogamy. The offspring resulting from this partnership are assigned to one of the genders with a 0.5 probability. The reproduction probability of the offspring is given by the average values of the two parents. The offspring is placed into a randomly chosen empty spot among the eight nearest neighbors of the parent that was originally picked for reproduction. If no empty spots exist within the immediate neighborhood, reproduction is not successful. Potential issues of mate preference for individuals with similar reproduction probabilities are not taken into account. Death occurs with a probability $D$, according to the same rules as described before.

As shown in figure 9, the outcomes remain robust. For relatively low population death rates, the reproduction probability can evolve towards reduced values until population extinction occurs. For relatively high death rates, the reproduction probability can evolve towards maximal values. For intermediate death rates, the population can fluctuate around an intermediate reproduction probability.

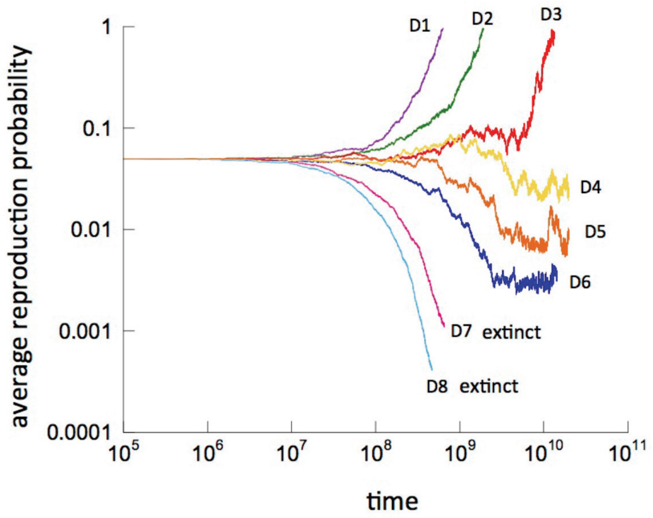

Figure 9: Outcomes of ABM4 with continuous reproduction strategies, cultural evolution, and sexual reproduction. Simulations are in principle the same as those presented in Figure 5 of the main text, but using the model with sexual reproduction. Assumed death rates decrease from D1 to D8. Results remain robust. Parameters were chosen as follows. Death rates are given by $D1 = 0.002, D2 = 0.001, D3 = 7.5 \times 10^{-4}, D4 = 7 \times 10^{-4}, D5 = 6.5 \times 10^{-4}, D6 = 5 \times 10^{-4}, D7 = 2.5 \times 10^{-4}, D8 = 1 \times 10^{-4}$. The reproduction probability of the individuals, R, was allowed to evolve, starting from R=0.05 among all individuals. The population death rate, $D$, is indicated in the graphs. $u = 0.1, M = 0.9; PC = 0.0003; Q = 0.965; G = 2\%$.

### 5 Model extensions – future work

Some processes in the more complex versions of the models considered here could also be formulated in slightly different ways. In ABM3 and ABM4, cultural transmission involves the calculation of the weighted average reproduction rate among individuals within the immediate neighborhood. The assumption was made that individuals with a faster reproduction rate than the agent under consideration count less in this process, irrespective of the magnitude of this difference. Alternatively, it could be assumed that the reduced weight is proportional to the difference in reproduction rates, thus taking into account the distance in social hierarchies. While it seems reasonable to assume that economically more successful individuals carry more weight in cultural transmission than individuals who are less successful, the details of this are not well understood. We note that results reported here depend on the assumption that individuals with lower reproduction rates carry more social weight, an assumption that has also been made in previous modeling work [1]. Another example of uncertainties in model construction is the formulation of the sexual reproduction model. We assumed monogamy, but made some obvious simplifications, as explained in the Results section. There are different assumptions that can be made in models that describe sexual reproduction, but the most important feature in the current context is that the reproduction rate of the offspring is not simply a copy of one of the parents, but represents the average of the two parents. This provides an additional mechanism of cultural change. Finally, only two types of communication networks have been considered in the agent based models here, the one where individuals interact with everyone else in the population, and the one where only interactions among nearest neighbors are allowed. A large variety of more realistic, random communication networks can be constructed, but we do not expect the results to differ from the ones obtained from the two extreme cases of networks considered here.
